# Ant abaecin-2 is a context-dependent copper-binding effector that can be either inhibitory or protective

**DOI:** 10.64898/2026.04.18.719391

**Authors:** Caroline M. Donaghy, Helena Heyer-Gray, Charlotte O’Hern, Michael Ibrahim, Bryant Perez-Torres, Nichali Bogues, Andrei T. Alexandrescu, Karrera Djoko, Jonathan L. Klassen, Alfredo M. Angeles-Boza

## Abstract

Host defense peptides (HDPs) are important components of the innate immune system that are used to combat pathogens and often rely on metal binding for their function. However, controlling trace nutrients such as transition metals may have other roles in host-symbiont interactions beyond poisoning harmful pathogens. This study characterizes the evolution, structural properties, and biochemical activity of the novel hymenopteran HDP abaecin-2. In myrmicine ants such as the fungus-growing tribe Attini, abaecin-2 has evolved to include an *A*mino-*T*erminal *C*u(II) and *N*i(II)-binding (ATCUN) motif, which we hypothesize may bind copper, a trace nutrient that is enriched in attine ant colonies. Combined results from mass spectrometry, competitive binding assays, circular dichroism, and NMR indicate that the abaecin-2 peptide lacks a defined secondary structure and can associate with up to 2 Cu(II) ions, one strongly bound at the ATCUN motif and another weakly bound, likely at a conserved histidine residue. Despite its copper-binding activity, abaecin-2 alone does not exhibit antibacterial activity against *Escherichia coli* or *Bacillus subtilis* (models for bacteria that live in ant fungus gardens). However, it synergizes with a model pore-forming peptide cecropin A to inhibit the growth of *E. coli*, similar to the related peptide abaecin-1. The copper-binding activity conferred by the ATCUN motif also protects copper-sensitive *E. coli* from excess copper toxicity. The dual, context-dependent inhibitory and protective roles we propose for abaecin-2 indicate that this previously under-characterized HDP may be used by attine ants to regulate both harmful and beneficial symbionts.

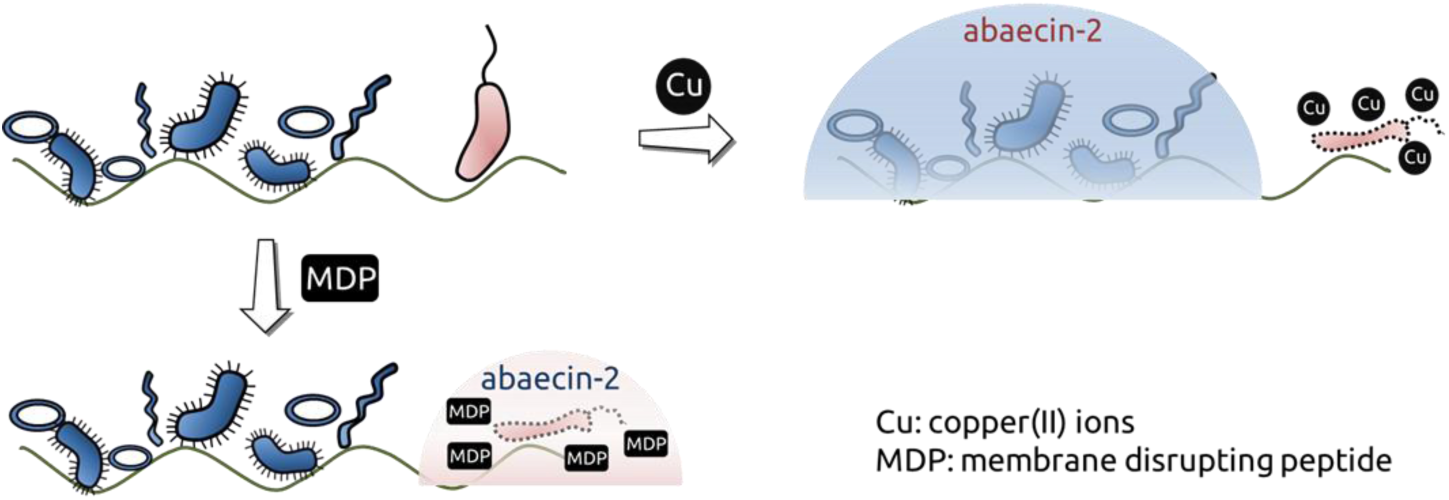

## Introduction

Molecules that control microbial populations within complex host-microbe symbioses using mechanisms other than direct toxicity have been underexplored. Instead, most studies of host defense molecules have centered on traditional antimicrobial agents, such as host defense peptides (HDPs). These are essential components of the innate immune system in all organisms, especially those that lack an adaptive immune system, including all insects^1^. Although many HDPs have diverse metal-binding motifs and display diverse metal-dependent mechanisms, the role of the metal is typically framed in the context of bacterial killing^2,3^.

How hosts control their microbial symbionts using host defense molecules is poorly understood, especially in multipartite mutualisms where hosts must manage both beneficial and pathogenic microbes. Such systems require resource management by the host, where specific nutrients, such as trace nutrient metal ions, can be either deployed at toxic concentrations to eliminate harmful pathogens or used to regulate beneficial symbiont growth^4^.

Fungus-growing ants (tribe Attini), for instance, host a complex symbiosis involving a cultivar fungus that provides the ants with food, fungal pathogens that infect either the ants or the cultivar fungus, and various beneficial bacteria^5^. Some fungus-growing ant colonies (*e.g.*, from the leaf-cutting genus *Atta*) form very large colonies containing millions of ants in hundreds to thousands of underground fungus garden chambers, and process huge amounts of plant material each day^6^. To thrive, these ants must not only exclude pathogens but also support the growth of beneficial symbionts. Fungus-growing ants support beneficial symbionts in part *via* nutrient provisioning, most notably to their cultivar fungus^7,8^. How nutrient provisioning helps control pathogens in the fungus-growing ant symbiosis is currently unknown, though the spatial distribution of nutrients is important^9^. The complexity of this mutualism likely necessitates host control over nutrient distribution to multiple symbionts, including *via* specific effector molecules.

Previous work indicates that copper ions may be a key regulatory nutrient in fungus-growing ant symbioses, because the leaf-cutting ant *Atta laevigata* selectively forages for copper-rich leaves^10^. Copper is a crucial nutrient for the ants, but its abundance must be carefully balanced so that it is tolerated by the cultivar fungus^7^. Other symbionts (*e.g.*, the gut bacterial symbionts *Mesoplasma* and *Spiroplasma*) lack recognizable genes that might protect them against copper toxicity^11^. A better understanding of host-derived chemical effectors capable of locally regulating copper availability, such as copper-binding peptides, may reveal how host-symbiont relationships are mediated by reconciling the different copper requirements of beneficial symbionts while excluding pathogens.

Insects frequently use HDPs to defend against microbial pathogens. For example, the HDP microplusin, isolated from the tick *Rhipicephalus microplus*, binds copper and exerts an antimicrobial effect by copper sequesteration^12^. Ants possess a suite of HDPs that includes abaecins, which are proline-rich peptides found in many hymenopteran insects^13^. Although the widely studied type 1 abaecins (abaecin-1) are generally considered weak antimicrobials that may require pairing with other peptides to increase their activity^14–16^, several ant species possess a second, distinct peptide called abaecin-2^13^. Unlike abaecin-1, many abaecin-2 peptides contain an <u>A</u>mino-<u>T</u>erminal <u>Cu</u>(II) and <u>Ni</u>(II) (ATCUN) binding motif^2^, which consists of a free terminal α-amino group and a histidine in the third position that together coordinate copper ions in a square-planar geometry^17,18^. The potential for two contrasting activities, *i.e.,* killing harmful symbionts (as an antimicrobial agent) *vs.* supporting beneficial symbionts (by mediating metal availability) makes such peptides ideal candidates for host-microbiome modulation.

In this work, we describe the interactions between copper and abaecin-2 from multiple fungus-growing ant species. Our phylogenetic analyses suggest that abaecin-2 emerged from an ancient gene duplication event in a hymenopteran ancestor and subsequently evolved to include an ATCUN motif in myrmicine ants, including all surveyed fungus-growing ants. We show that copper accumulates in the fungus garden and worker ants of two different species studied by our labs, *Atta texana* and *Trachymyrmex septentrionalis.* We further establish that abaecin-2 is a largely disordered peptide that has a high affinity for Cu(II). This high affinity can be partially attributed to the presence of the ATCUN motif. When combined with a membrane-disrupting peptide, abaecin-2 acts as an antibacterial agent, potentially by delivering bound copper to an unknown intracellular target. However, abaecin-2 can also bind excess copper and protect copper-sensitive microbes from copper toxicity. Taken together, these results indicate that abaecin-2 serves dual roles: 1) acting as an antibacterial agent when combined with a membrane-disrupting peptide; and 2) preventing copper toxicity in copper-sensitive symbiotic bacteria.

## Materials and Methods

### Phylogenetic analysis of abaecin-2

Insect abaecin-1 and -2 sequences were identified by performing two Position-Specific Iterated BLAST (PSI-BLAST)^19^ searches of the NCBI non-redundant protein sequences database. An *A. columbica* abaecin-2 sequence (Table S1) was used as the initial query sequence. A putative *Pogonomyrmex barbatus* abaecin-2 sequence returned in the initial PSI-BLAST search (Table S1) was used as the query sequence in the second search. Both searches were run until no additional unique sequences were returned. The unique sequences recovered by both searches were combined, and signal peptide sequences were removed as previously described^13^. Trimmed abaecin-1 and abaecin-2 sequences were aligned using MUSCLE with default parameters in the Molecular Evolutionary Genetics Analysis (MEGA) v. 11.0.13 package^20^. A maximum-likelihood (ML) tree was estimated using MEGA as described by Hall^21^ using the JTT+G model with 500 bootstrap replicates.

### Evolutionary history of the ATCUN motif in ant abaecin-2

To identify potential unannotated abaecin-2 sequences in ant genomic DNA, we performed translated BLAST (tBLASTn) searches against the NCBI whole genome shotgun contigs (WGS) database using 9 ant abaecin-2 sequences (Table S1) as query sequences and limiting the search results to sequences from Formicidae (taxid:36668). *A. texana* and *A. colombica* sequences were previously published, and the other sequences used were identified as putative abaecin-2 sequences via PSI-BLAST searches as described above. The unique nucleotide sequences returned by each query were translated to protein sequences using the batch translation tool at https://www.bioinformatics.org/sms2/translate.html^22^. Signal peptide sequences that preceded the ATCUN motif were removed in all translated protein sequences as described above. We acknowledge that the 3’ ends of some peptides predicted from our WGS searches may be inaccurate due to the presence of short 3’ exons that cannot be easily predicted using tBLASTn.

All ant abaecin-2 sequences identified using tBLASTn searches and 4 wasp abaecin-2 sequences identified by the initial PSI-BLAST search (*Vespula pensylvanica*, *Vespula germanica*, *Vespula vulgaris*, and *Mischocyttarus mexicanus*) were aligned using MUSCLE in MEGA as described above. Substitution models with the lowest Bayesian Information Criterion (BIC) score were chosen to estimate trees for each dataset. An ML tree of all sequences was estimated using the JTT+G model with 500 bootstrap replicates. A second ML tree containing one representative abaecin-2 sequence per ant genus and using the *M. mexicanus* and *V. pensylvanica* sequences as outgroups was estimated using the cpREV model with 500 bootstrap replicates. Both datasets also included a published *S. invicta* abaecin-2 sequence^13^. Alignments were visualized using Jalview^23^.

### Sample collection and ICP-MS analysis

Soil, fungus garden, and worker ants were collected in May 2021 following Permit WL-Research 2016-10 issued by the Louisiana Department of Wildlife and Fisheries. *A. texana* colonies were sampled in the Clear Creek Wildlife Management Area, Louisiana, USA. *T. septentrionalis* colonies were sampled in the Clear Creek Wildlife Management Area and Alexandria State Forest, Louisiana, USA (Table S3). Samples were placed in sterile 15 mL Falcon tubes using sterilized stainless-steel instruments. All samples were flash-frozen on dry ice and transported back to the University of Connecticut (Storrs, Connecticut, USA) for analysis.

The copper contents of all samples were measured using an Inductively Coupled Plasma – Mass Spectrometer (ICP-MS; Perkin Elmer/DRC-e, Perkin Elmer, Shelton, CT, USA). The ants and fungus garden were weighed individually, homogenized, and portioned to a final weight of 0.1-0.5 g. Ants, fungus garden, and plants were extracted following the EPA 200.3 method^24^. Soil was digested following the EPA 3050B method^25^ and had a final weight of 0.5 ± 0.1 g. The quality controls used for these experiments included the inclusion of duplicate samples, laboratory fortified matrices, laboratory reagent blanks, and laboratory fortified blanks. The standard reference materials used were DOLT-5 for biological tissues and PACS-3 for soil samples.

### Synthesis of abaecin-2 peptides for characterization

The abaecin-2 peptides used for this study were either derived from the published genome sequences of *A. texana* or *A. colombica* ants. Each abaecin-2, their respective H3A mutants, magainin-2, and cecropin A peptides were synthesized using a Syro I peptide synthesizer from Biotage (Charlotte, NC, USA). The synthesis of the peptides used fluorenylmethyloxycarbonyl (Fmoc) chemistry on Low-Loading (LL) Rink amide resin (ChemPep Inc, Wellington, FL, USA). After cleavage, the peptides were purified by high-performance liquid chromatography (HPLC; Thermo Scientific Dionex UltiMate 3000 HPLC, Waltham, MA USA) using a C18 semipreparative column (Jupiter® C18 column, Phenomenex, Torrance, CA, USA), and their masses were confirmed via electrospray ionization mass spectrometry in positive ion mode (4000 QTRAP, AB Sciex, Marlborough, MA, USA). Peptide purity was determined using reverse-phase HPLC with a C18 column (Jupiter® C18 column, Phenomenex, Torrance, CA, USA). The fractions containing the purified peptide were lyophilized overnight and dissolved in nanopure water. Each peptide was quantified via UV−vis spectrophotometry using molar extinction coefficients for the amino acid residues and the peptide bond as determined by Kuipers and Gruppen^26^. All peptides were lyophilized after synthesis, and re-suspended and re-quantified before any new experiment.

### Copper binding experiments

To measure copper binding by mass spectrometry, a 10 mM stock of Cu(II) was prepared from CuCl_2_ in nanopure water. Peptides were prepared to a final concentration of 100 μM in 500 μL of nanopure water. Peptides exposed to copper were prepared as solutions of 1:5 peptide:Cu(II) to a final volume of 500 μL and incubated for 30-60 minutes before injection. Samples were analyzed on a 4000-Q TRAP LC/MS/MS system by Applied Biosystems (Waltham, MA, USA). Data were deconvoluted from 600 to 1400 m/z using UniDec^27^ (https://unidec.chem.ox.ac.uk/) to obtain masses at 1 Da resolution.

For the desalting experiments, *apo-*peptides (50 µM each) were incubated with excess CuCl_2_ (>100 µM) for 15 min in morpholinepropanesulfonic acid (MOPS, Sigma-Aldrich, Burlington, MA, USA) buffer (50 mM, 100 mM NaCl, pH 7.4). This mixture was eluted through a PD-10 desalting column (Cytiva, Marlborough, MA, USA). Peptide concentrations in each eluted fraction were determined using a QuantiPro BCA Assay Kit (Merck, Rahway, NJ, USA). The copper concentrations in the same fractions were determined using ICP-MS at the Metallomics Facility, Department of Biosciences, Durham University.

Cu(II) binding affinities were estimated by competition with fluorometric Cu(II)-binding probes DP2 (log *K*_D(Cu)_ = -10.1), DP3 (log *K*_D(Cu)_ = -12.3), and DP4 (log *K*_D(Cu)_ = -14.1) as previously described^28^. A master solution containing an *apo*-peptide (28.8 or 115.2 µM) and a probe (18 or 36 µM) was prepared in MOPS buffer (50 mM, 100 mM NaCl, pH 7.4). A series of CuCl_2_ stock solutions (0 – 120 µM) was prepared separately in water. Exactly 180 µl of the master solution was added to 20 µl of each CuCl_2_ stock solution in black µClear® microplates (Greiner, Kremsmünster, Austria). After 10 min, fluorescence emission intensities at 550 nm following excitation at 350 nm were recorded using a Synergy H4 plate reader (BioTek, Winooski, VT, USA). Fluorescence values (*F*_x_) were normalised such that the maximum (*F*_max_, at 0 µM Cu(II)) and minimum (*F*_min_, at excess Cu(II)) values represented 100% and 0% *apo*-probe, respectively. The ratios of *apo*-probe to total probe concentrations were plotted against final Cu(II) concentrations. To obtain *K*_D(Cu)_ values for each peptide, the resulting competition curves were fitted using DynaFit (BioKin Ltd., Watertown, MA, USA) and a script describing the relevant Cu(II)-peptide and Cu(II)-probe binding equilibria (Supplementary code S1).

### Circular Dichroism (CD) spectroscopy

Abaecin-2 WT and H3A mutant peptides from *A. texana* and *A. columbica* were each diluted to a final concentration of 100 μM in 350 μL of 10 mM potassium phosphate buffer (KPB) at a pH of 7.4. CD spectra of these peptides were collected either in KPB or in a 1:2 ratio of peptide:Cu(II) with 200 μM of Cu(II) added to either solution. Each sample was run using an Applied Photophysics Chirascan V100 Spectrometer (Surrey, UK) in a 0.1-cm quartz cuvette at room temperature. All samples were run with at least two replicates, and spectra were collected at 1.0 nm intervals from 250 to 190 nm. The bandwidth was 3 nm. When metal ions were added to the peptide solutions, these samples were incubated for 45 to 60 min before analysis. All data were input into the BeStSel software^29^ (http://bestsel.elte.hu/index.php) for secondary structure analysis.

### NMR spectroscopy

NMR assignments were obtained using a sample of 1 mM WT *A. texana* abaecin-2 in 90% H_2_O/10% D_2_O at pH 6.6. A temperature of 10 °C was used to reduce amide proton exchange with the solvent^30^. Residue-specific NMR assignments were determined from sequential walks^31^ using 70 ms mixing time TOCSY (pulse sequence *dipsi2gpph19*) and 250 ms mixing time NOESY (Nuclear Overhauser Effect Spectroscopy; *noesygpph19*) spectra recorded on an 800 MHz Bruker (Billerica, MA, USA) Avance Neo spectrometer with a cryogenic probe. Additional data for spin-system assignments included multiplicity-edited ^1^H-^13^C HSQC (*hsqcedetgpsosp2.3*) spectra for both WT and H3A mutant abaecin-2, as well as 70 ms mix time TOCSY spectra recorded on 0.5 mM samples of WT and H3A in 99.96% D_2_O at pD 6.6 ± 0.2 and a temperature of 32 °C. NMR assignments for WT abaecin-2 peptides have been deposited in the BMRB under accession code 52634.

Random coil index values were calculated using the RCI server (https://www.randomcoilindex.ca)^32^. The binding of Cu(II) for WT and H3A variants of abaecin-2 on a per-residue basis was characterized using the loss of Hα-Cα NMR cross-peak volumes in the presence of the paramagnetic metal. NMR assignments and cross-peak volume measurements were done using the program CcpNmr Analysis v. 2.5.2^33^.

### Antimicrobial susceptibility assay

Antimicrobial activity was determined using the broth microdilution method^34^. One to two colonies of *E. coli* MG1655, *E. coli* JW 0473-3 (Δ*copA*), or *E. coli* JW 0013-4 (Δ*dnaK*) were added to MHB (MilliporeSigma, Burlington, MA, USA), unadjusted to maintain pH 7.4, or minimal media (1xM9 salts, 0.2% glucose) that was adjusted to pH 7.4 using MOPS. These cultures were incubated until the mid-logarithmic phase was achieved, at an OD_600_ of 0.4−0.6. Bacterial cultures containing 1 × 10^6^ colony-forming units (CFU)/mL were then prepared in fresh media for the determination of values. Peptide stock solutions were prepared at 2 times the maximum concentration needed for the experiment, and 50 μL of 2-fold serial dilutions were made in a 96-well polypropylene plate (Greiner, Kremsmünster, Austria). After the serial dilutions, 50 μL of each bacterial suspension of 1 × 10^6^ CFU/mL was added, achieving a final concentration of 5 × 10^5^ CFU/mL. Plates were then incubated at 37 °C for 16-18 hrs, and the MIC was observed as the lowest concentration that visibly inhibited bacterial growth. The MIC values reported represent the mode of three or more trials.

A standard checkerboard assay^35–37^ was used to elucidate the antimicrobial activity of *A. texana* and *A. colombica* WT and H3A abaecin-2 peptides paired with the well-characterized pore-forming peptides cecropin A and magainin-2. Stock solutions of each peptide were prepared to 8× their expected MIC. 100 μL of the first peptide stock solution was added to the first 8 wells of the first column of a 96-well plate. The peptide was serially diluted 1:2 by moving 50 μL from each of the wells in the first column into the adjacent wells in the second column of the plate, which contained 50 μL of either MHB or M9 media. This serial dilution was repeated for columns 3-7 of the plate, leaving 50 μL of media in each well of column 8. The second peptide of interest was serially diluted in an additional 96-well plate by first adding 100 μL of stock solution to each of the 8 wells in row A of the plate, then transferring 50 μL of the undiluted stock solution from each well in the first row to the well below it, which contained 50 μL of either MHB or M9 media without additional Cu(II) supplementation. This process was repeated for rows C-G of the plate, leaving 50 μL of media in the wells in row H.

Once the initial peptide dilutions were completed, 50 μL of the peptide suspension in each well of the second plate was added to the matching wells. Bacteria of interest were grown in MHB or M9 media until mid-logarithmic growth (as determined by OD_600_) was reached. Cells were diluted to a concentration of 1 × 10^6^ CFU/mL, and 100 μL of this cell suspension was added to each well for a final concentration of 1 × 10^4^ CFU/mL in each well with peptides. Plates were incubated at 37 °C for 16-18 hours, then read visually to determine the well with the lowest concentration of both peptides that had no bacterial growth. The concentration of each peptide in these wells was used to calculate the Fraction Inhibitory Concentration (FIC) index as follows:

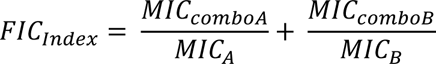

An MIC of 256 μM was used for abaecin-2 peptides to take a conservative approach in calculating synergy. FIC indices ≤ 0.5 indicate synergy between the two peptides being tested, while indices 0.5-4 indicate additive behavior, and indices > 4 indicate antagonistic behavior between the peptides^36–38^. Although synergy corresponds to FIC indices < 1 mathematically, we and others have suggested this more stringent standard to accommodate experimental uncertainties in the determination of MIC values^36^. FIC indices reported here are the mode of three trials that were all done in duplicate.

### LPS binding experiment

To determine whether affinity for LPS plays a role in its antibacterial activity, the LPS binding ability of WT and H3A *A. texana* abaecin-2 was assessed using the BODIPY-TR-cadaverine (BC) substitution assay^39^. LPS-BC master mix was prepared by incubating 3.5 μg/mL LPS (*E. coli* O111:B4) with 5 μM BC in 50 mM Tris buffer (pH 7.4) in the dark for 15 minutes. Antibiotics and antimicrobial peptides with known affinities for LPS (high affinity: polymyxin B, magainin 2, cecropin A; low affinity: streptomycin or vancomycin) were used as controls. Stock solutions of WT and H3A abaecin-2 and each control were prepared to a concentration of 64 μM and serially diluted 1:2 with equal volumes of LPS-BC master mix for final peptide/antibiotic concentrations of 32, 16, 8, 4, 2, 1, 0.5, or 0.25 μM. Fluorescence intensities using excitation and emission wavelengths of 580 and 620 nm, respectively, were measured using a Biotek Cytation 5 fluorescence spectrophotometer (Agilent Technologies, Santa Clara, CA) every 5 minutes for 45 minutes. All assays were repeated a total of three times.

### Bacterial rescue experiment

One to two colonies of *E. coli* MG1655 or *E. coli* JW 0473-3 (Δ*copA*) were inoculated into 1x M9 media and cultured overnight. These cultures were then washed three times with 1x M9 media by repeated centrifugation at 12,000 × g, removal of the supernatant, and resuspension in new 1x M9 media. After the cells were washed, they were resuspended in a 1:100 dilution of 1x M9 media and regrown to their logarithmic growth phase. These cells were diluted 1:100 in 1x M9 media to a final volume of 2.5 mL. A 10 mM solution of CuCl_2_ was prepared in sterile nanopure water and filtered through a Rapid-Flow 0.2 μm polyethersulfone filter (Nalgene, Rochester, NY, USA). This Cu(II) solution was used to prepare 0, 0.1, 0.5, 1, 5, and 10 μM dilutions that would be tested in the absence of peptides or with either 10 μM abaecin-2 or H3A abaecin-2 from *A. texana.* These solutions were incubated for 24 hours at 37 °C while shaking at 150 rpm. After incubation, the optical density at 600 nm was observed using a Varian Cary 50 Scan UV-Vis spectrophotometer (Agilent Technologies, Santa Clara, CA, USA).

## Results and Discussion

### Abaecin-2 arose from a gene duplication and occurs in many hymenopteran species

PSI-BLAST searches of the NCBI database (using query sequences in Table S1) returned 54 unique protein sequences (42 abaecin-1 and 12 abaecin-2) from 33 hymenopteran genera in 8 families (Figure 1). Phylogenetic tree reconstruction revealed two distinct clusters corresponding to abaecin-1 and abaecin-2, clearly separating the different forms of abaecin produced by the same species. Abaecin-1 and abaecin-2 are both present in ant and social wasp species, but only abaecin-1 is present in bees, solitary wasps, one sawfly (*Athalia rosae*), and one social wasp (*Vespa crabro*). This phylogenetic distribution suggests loss of abaecin-2 genes in at least some of these latter species, following a gene duplication in their shared ancestor.

**Figure 1.**
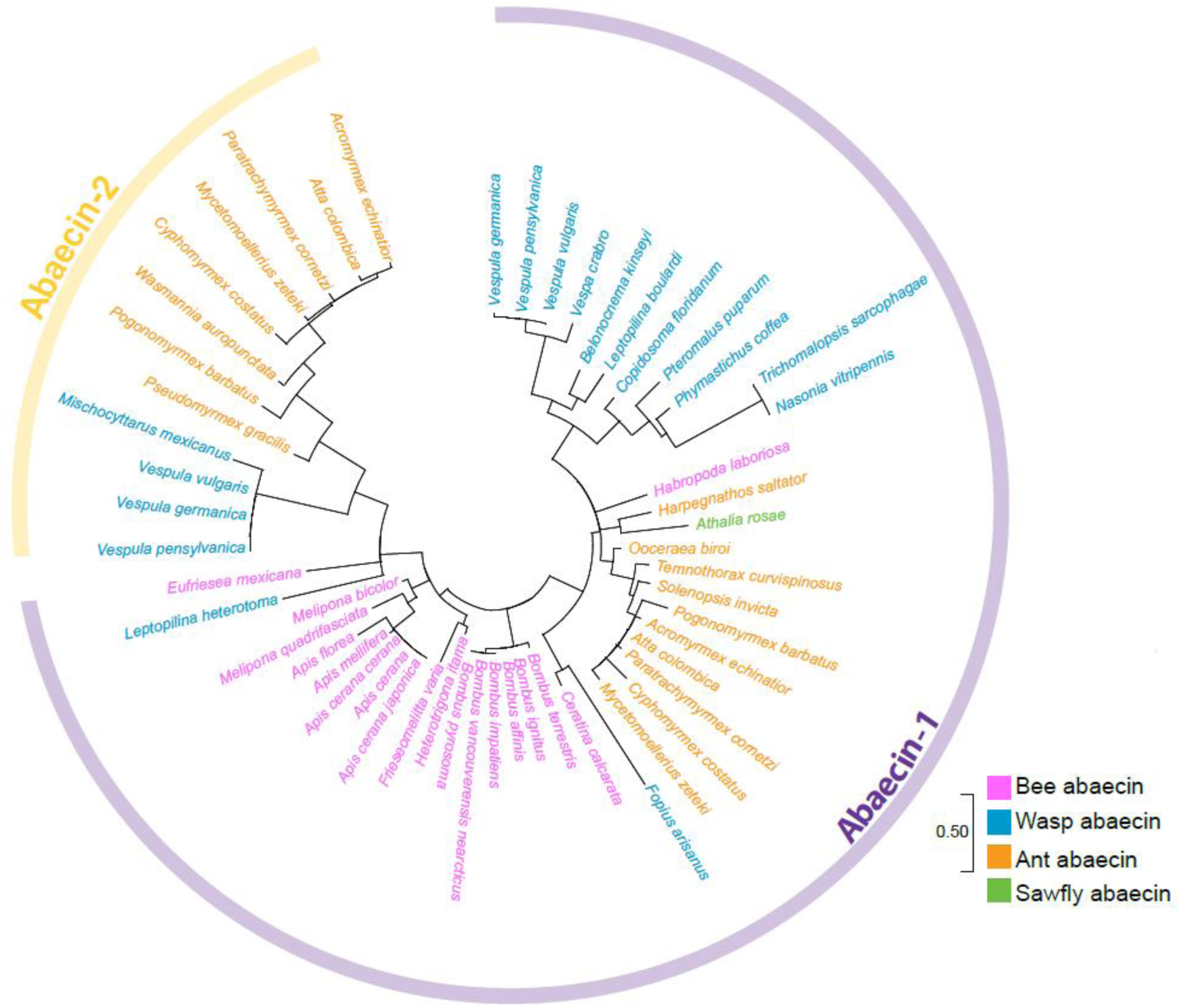
Distribution of abaecin peptides in hymenopteran species. Maximum likelihood tree of abaecin-1 (n = 42) and abaecin-2 (n = 12) protein sequences. Sequences cluster into two distinct clades, each corresponding to a different form of abaecin. The scale bar corresponds to the number of amino acid substitutions per site.

### Ants evolved an ATCUN motif in their abaecin-2 peptides

tBLAST searches of the NCBI database (using query sequences in Table S1) returned 58 unique ant abaecin-2 sequences, from which 25 randomly selected representatives from different genera were used to estimate a maximum likelihood (ML) tree that also included a previously published *Solenopsis invicta* abaecin-2 sequence (Figure 2). Additional ML trees were also estimated from all 58 ant abaecin-2 protein and nucleotide sequences (Figure S1, S2). Rooting these ant abaecin-2 trees using their wasp homologs suggests that the ATCUN motif (specifically the amino acid sequence DTH) gradually evolved from an ancestral peptide that did not contain this motif, having evolved in only one of the two major clades of ant abaecin-2 sequences. This clade contains sequences from both fungus-growing ants (tribe Attini) and non-fungus-growing ants from the tribes Stenammini, Solenopsidini, and Crematogastrini. A second His residue at position 9 is present in all abaecin-2 sequences with a DTH motif, while a third His residue at position 36 is highly conserved in attine ants but not in other species.

**Figure 2.**
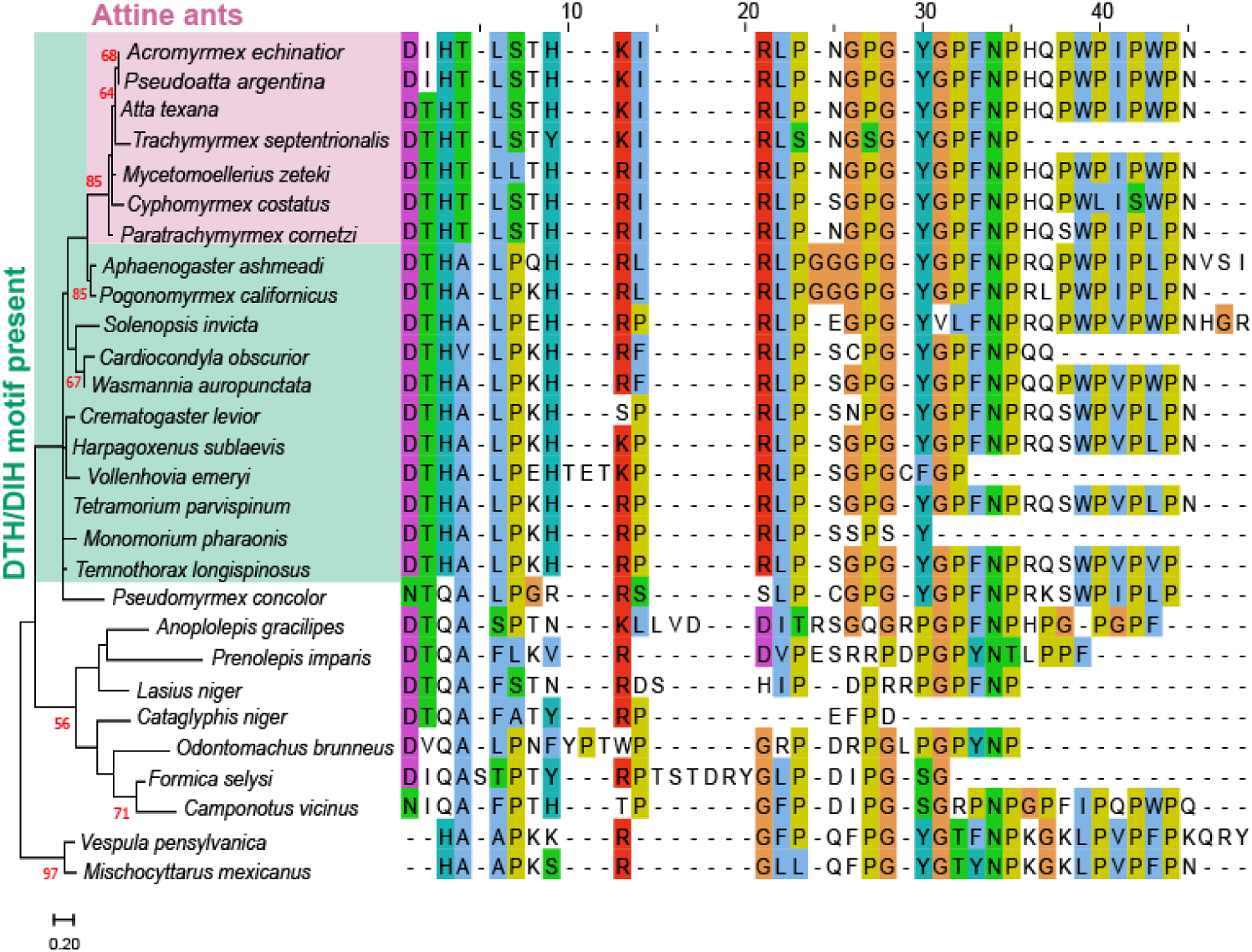
Evolution of the ATCUN motif in ants. Maximum likelihood tree of 25 representative ant abaecin-2 sequences. The DTH version of the ATCUN motif (tree: green box) emerged from an ancestral version of the peptide and is conserved in most attine ants (tree: pink box). Some sequences may be incomplete due to the difficulty of predicting very short 3’ exons based on sequence homology alone. The scale bar corresponds to the number of amino acid substitutions per site.

Although the poor bootstrap support for large portions of the tree makes it difficult to definitively determine the origin of the DTH motif, the most parsimonious reconstruction is that the ancestral ant sequences had both D and T (but not H) residues, with the H residue being acquired during the early evolution of the ant subfamily Myrmicinae. The DTH motif may have subsequently been lost in *Pseudomyrmex* abaecin-2 sequences, and an additional version of the ATCUN motif (DIH) evolved more recently among attine ants. Although the ATCUN motif emerged before the mutualistic relationship between fungus-growing ants and their cultivar, the conservation of this motif and the emergence of an additional conserved His residue in abaecin-2 peptides from fungus-growing ants suggest that the copper-binding properties of this peptide may play a key role in symbiont control by regulating copper availability.

### Copper is differentially enriched in fungus-growing ant colonies

To determine whether copper is selectively enriched in fungus-growing ant colonies, as has been previously shown for the fungus-growing ant *A. laevigata*^10^, we assessed the amounts of copper in *A. texana* and *T. septentrionalis* nests using Inductively Coupled Plasma-Mass Spectrometry (ICP-MS). Worker ants, fungus gardens, and soil surrounding nest chambers were collected in the field before analysis in the lab.

In *T. septentrionalis* colonies, copper contents (parts per billion, ppb) differed between the ants, their fungus garden, and the surrounding soil (Figure 3; 1-way ANOVA, F = 81.99, df = 2,37, p <<0.001). The copper content in worker ants was 127% higher than in the fungus garden (p = 0.006) and 730% higher than the surrounding soil (p < 0.001). The fungus garden was also enriched (226% higher) in copper compared to the surrounding soils (p << 0.001). In micromolar units, these concentrations correspond to approximately 30 µM in soil, 80 µM in the fungus garden, and roughly 200 µM in the ants.

**Figure 3.**
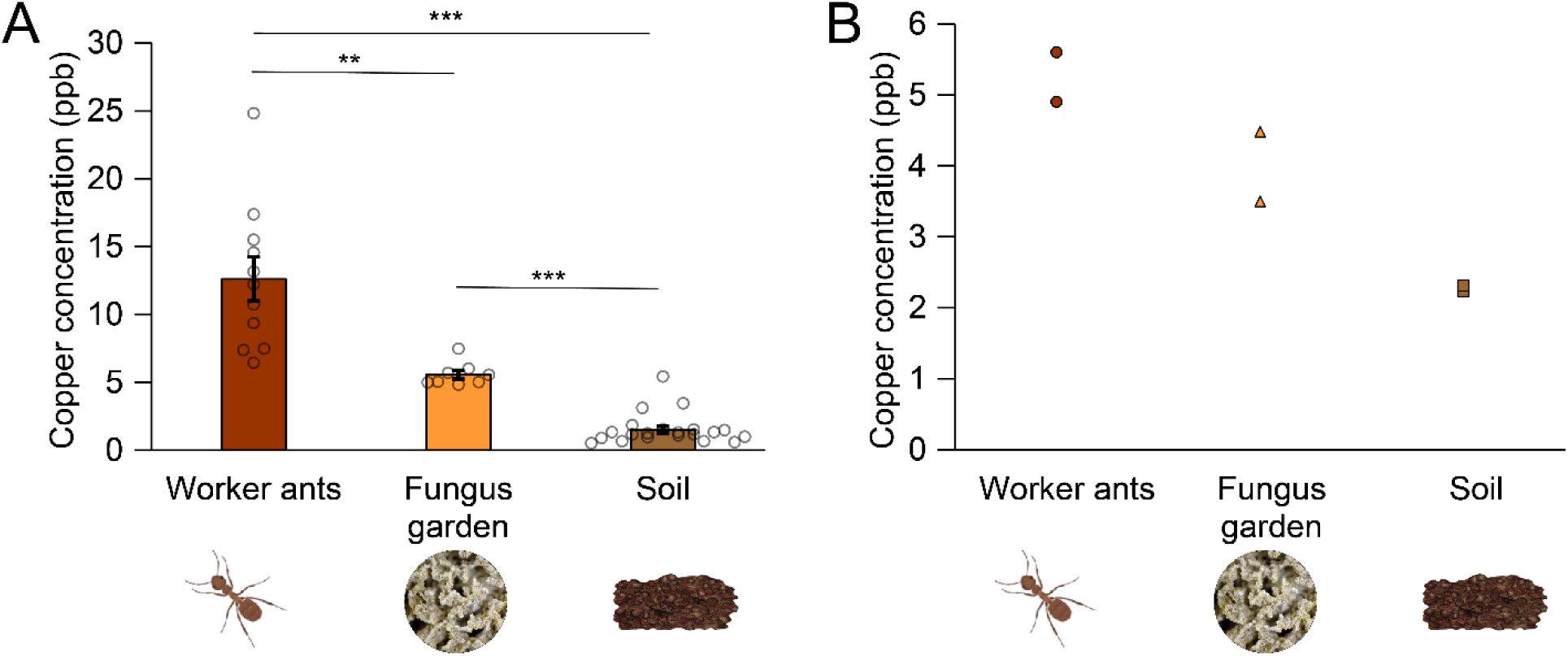
Relative abundance of Cu(II) ions in *T. septentrionalis* and *A. texana* colonies. Mean (± SE) Cu(II) contents (ppb) measured in worker ants, fungus gardens, and soil using ICP-MS. **A)** Cu(II) ion contents differed significantly between worker ants (n = 11), fungus gardens (n = 8), and soil (n = 21) from 11 *T. septentrionalis* nests (p << 0.001). All sample types were significantly different from each other according to a Tukey’s HSD test. **B)** Cu(II) ion contents varied significantly based on sample type from 2 *A. texana* colonies (p < 0.05). Worker ant (n = 2) and soil (n = 2) samples, and fungus gardens n = 2) and soil samples, differed significantly from each other according to a Tukey’s HSD test; *: p < 0.05, **: p < 0.01, ***: p< 0.001. Note that the y-axis scales are different for each panel.

In *A. texana* colonies, copper content differed depending on sample type (Figure 3; 1-way ANOVA, F = 27.18, df = 2,3, p = 0.012). The copper content of worker ants was 131% higher than the surrounding soil (p < 0.05), and the copper content of the fungus garden was 76% higher than the surrounding soil (p < 0.05). However, the copper content of worker ants and the fungus gardens did not differ from each other (p = 0.179). In both species, worker ants had the highest copper contents, followed by the fungus garden (Figure 3). Note that all of these are bulk measurements that combine multiple tissues or microhabitats, and so local copper concentrations (i.e., those experienced by microbial symbionts) may be considerably higher or lower than our measurements. The accumulation of copper in the ants and fungus garden suggests that it may be a key nutrient in this system, similar to what has been suggested for a related species^10^. However, even if copper is a necessary nutrient for the ants, copper levels are likely still regulated within ant colonies due to its potential toxic effects on symbionts at high levels, a pattern that has been observed for other trace minerals^7^.

### Abaecin-2 binds copper ions

The accumulation of copper in ant colonies implies a need for the ants and their microbial symbionts to manage this metal. Peptides containing an ATCUN motif are known to bind Cu(II) ions with high affinities^17,40^. We therefore characterized the Cu(II)-binding properties of abaecin-2 as a candidate mechanism for copper management^17,40^. To determine whether copper binding by abaecin-2 also depended on the ATCUN motif, we examined the binding of Cu(II) to both wild-type (WT) abaecin-2 and a synthetic H3A mutant lacking the ATCUN histidine at position 3, which is expected to have lower affinity for this ion^41^. Only data for abaecin-2 from *A. texana* is shown here because we could not accurately predict the 3’-end of the *T. septentrionalis* abaecin-2 sequence, owing to the difficulty of detecting its putative short 3’ exon based on nucleotide homology alone. Some experiments were also repeated with abaecin-2 from *A. columbica* (Figure S4, Figure S5, Table S4); these were entirely consistent with the experiments conducted using *A. texana* abaecin-2.

To characterize abaecin-2’s ability to bind Cu(II), both WT and H3A mutant peptides from *A. texana* were incubated with 5 molar equivalents of Cu(II) at pH 7, and the mixtures were separated on desalting columns and analyzed using native high-resolution electrospray ionization mass spectrometry (HR-ESI-MS; Figure S3). The resulting deconvoluted spectra suggest that both the WT and H3A abaecin-2 peptides associate with up to two Cu(II) ions each (Figure S3). Upon removal of excess and weakly bound Cu(II) using a desalting column, 0.9 ± 0.2 molar equivalents of Cu(II) ions remained bound to the WT peptide (Figure 4, Figure S4). This suggests that only one of the two Cu(II) ions detected in the HR-ESI-MS experiment was bound with high affinity, likely at the ATCUN motif. Consistent with this suggestion, the H3A variant co-eluted with only 0.4 ± 0.1 molar equivalent of Cu(II) (Figure 4, Figure S4). This sub-stoichiometric value indicates partial dissociation of weakly-bound Cu(II) from the H3A abaecin-2 variant on the desalting column.

**Figure 4.**
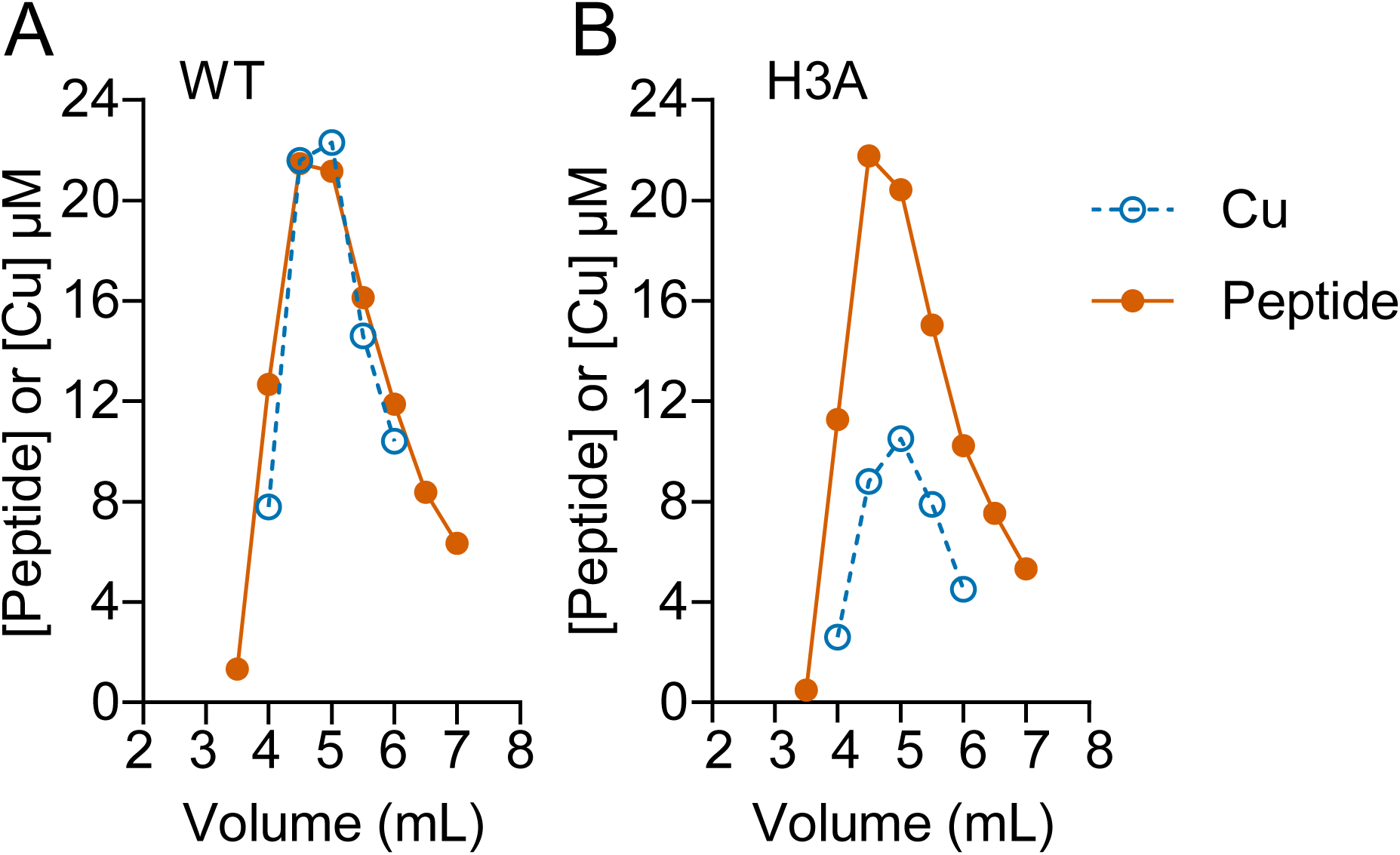
Copper recovery from abaecin-2 after desalting. Elution of WT (A) and H3A (B) abaecin-2 peptides from *A. texana* on a desalting column. Data points show the concentrations of peptide (filled circles, ●) and Cu(II) ions (open circles, ○) in each fraction. Data are from one representative of three independent experiments.

To estimate Cu(II)-binding affinities, WT and H3A mutant abaecin-2 peptides from *A. texana* were competed against fluorometric Cu(II) probes with known affinities.^42^ The resulting competition curves (Figure 5, Figure S5) indicated that WT abaecin-2 bound Cu(II) with a femtomolar affinity (*K_D_* = 4.4 ± 0.5 x 10^-14^ M), consistent with the expected high affinity of an ATCUN motif^17,40^. As expected, loss of His3 in the ATCUN motif of the H3A peptide led to a 10,000-fold reduction in affinity (*K_D_* = 4.5 ± 1.0 x 10^-10^ M).

**Figure 5.**
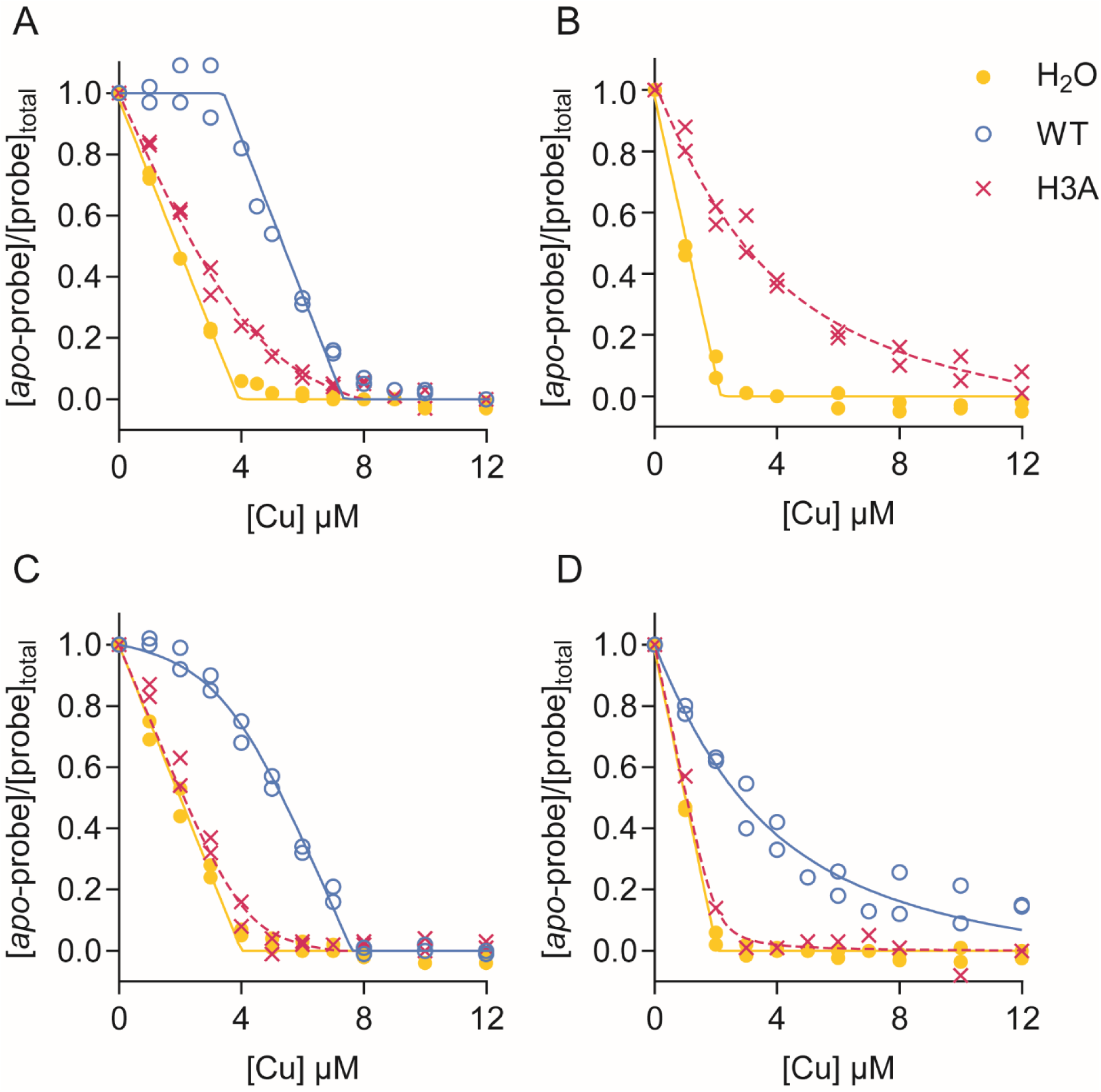
Cu(II) binding affinity of *A. texana* abaecin-2. The WT (○, solid lines) or H3A (x, dashed lines) peptides were competed against the fluorescence probe DP2, DP3, or DP4. Data points (n = 2) show the resulting competition curves when Cu(II) was added to mixtures of (A) 4 µM DP2 and 3.2 µM peptide, (B) 2 µM DP2 and 12.8 µM peptide, (C) 4 µM DP3 and 3.2 µM peptide, or (D) 2 µM DP4 and peptide. Control curves without any peptide are also shown (●, solid lines),

### Abaecin-2 adopts a disordered structure

The proline-rich sequence of abaecin-2 suggests that it should have a disordered structure, as previously estimated for *A. mellifera* abaecin-1^43^. Based on analysis of circular dichroism (CD) spectra using the Beta Structure Selection (BeStSel) software^29^, the secondary structures of the *A. texana* WT and H3A abaecin-2 peptides, with and without copper (Figure 6A), are dominated by “other” and turn structure (Figure 6B)^29^. When exposed to copper, the helical propensity of both the WT peptide and H3A mutant did not change significantly. The abaecin-2 structure predicted by AlphaFold 3 (AF3)^44^ in the presence of a single Cu(II) ion also has little regular hydrogen-bonded secondary structure. Although the AF3 pLDDT scores indicated a confident prediction (90> pLDDT > 70), the top 5 models generated by AF3 showed poor convergence with RMSD values after best-fit alignment of between 4 and 9 Å. The AF3 results thus also suggest that abaecin-2 lacks a unique folded structure.

**Figure 6.**
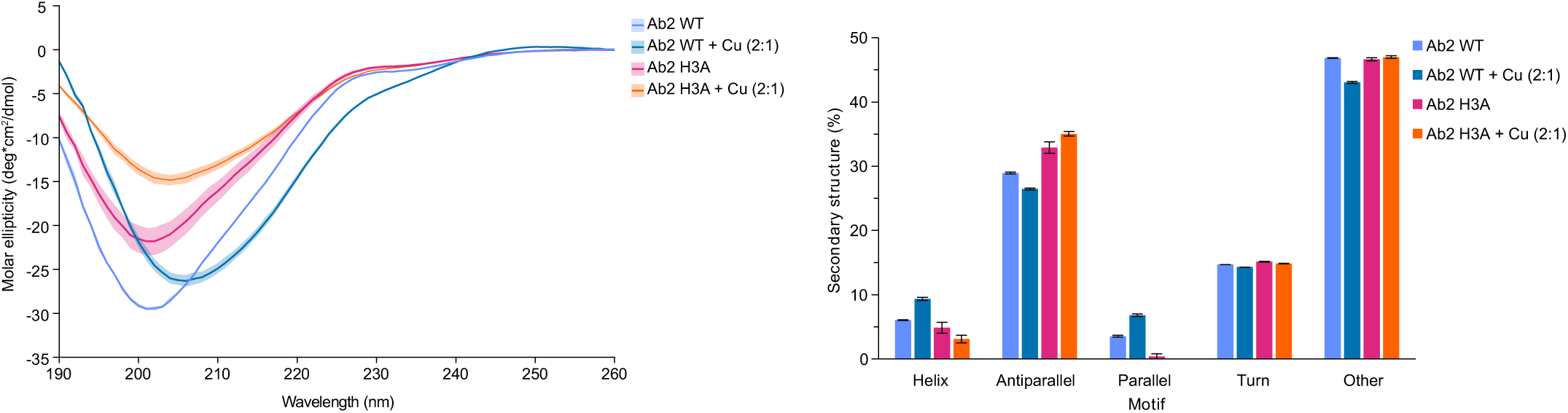
Secondary structure analysis of *A. texana* abaecin-2 peptides. (A) The CD spectra for 100 µM WT or H3A abaecin-2 peptides with or without 2 molar equivalents of Cu(II) in potassium phosphate buffer (KPB). (B) Secondary structure predictions show no major changes in the structure percentages of either peptide after adding Cu(II).

### NMR supports Cu(II) binding by histidines and the ATCUN motif in an otherwise unfolded peptide

To further characterize the structural and metal-binding properties of abaecin-2, we studied Cu(II) binding to the *A. texana* WT and H3A peptides using paramagnetic NMR line broadening (Figures S6-S8), in particular using multiplicity-edited ^1^H-^13^C HSQC spectra obtained at natural isotope abundance (Figure 7). Quantitative residue-resolution level information on paramagnetic broadening induced by Cu(II) binding^45^ was obtained from analysis of cross-peak volumes in the fingerprint Hα-Cα region of the ^1^H-^13^C HSQC spectrum (Figure 7B). Each NMR cross-peak was assigned as described in the Methods section. Resonances for the H3A mutant were assigned based on differences in the spectra compared to the WT (Figure S9).

**Figure 7.**
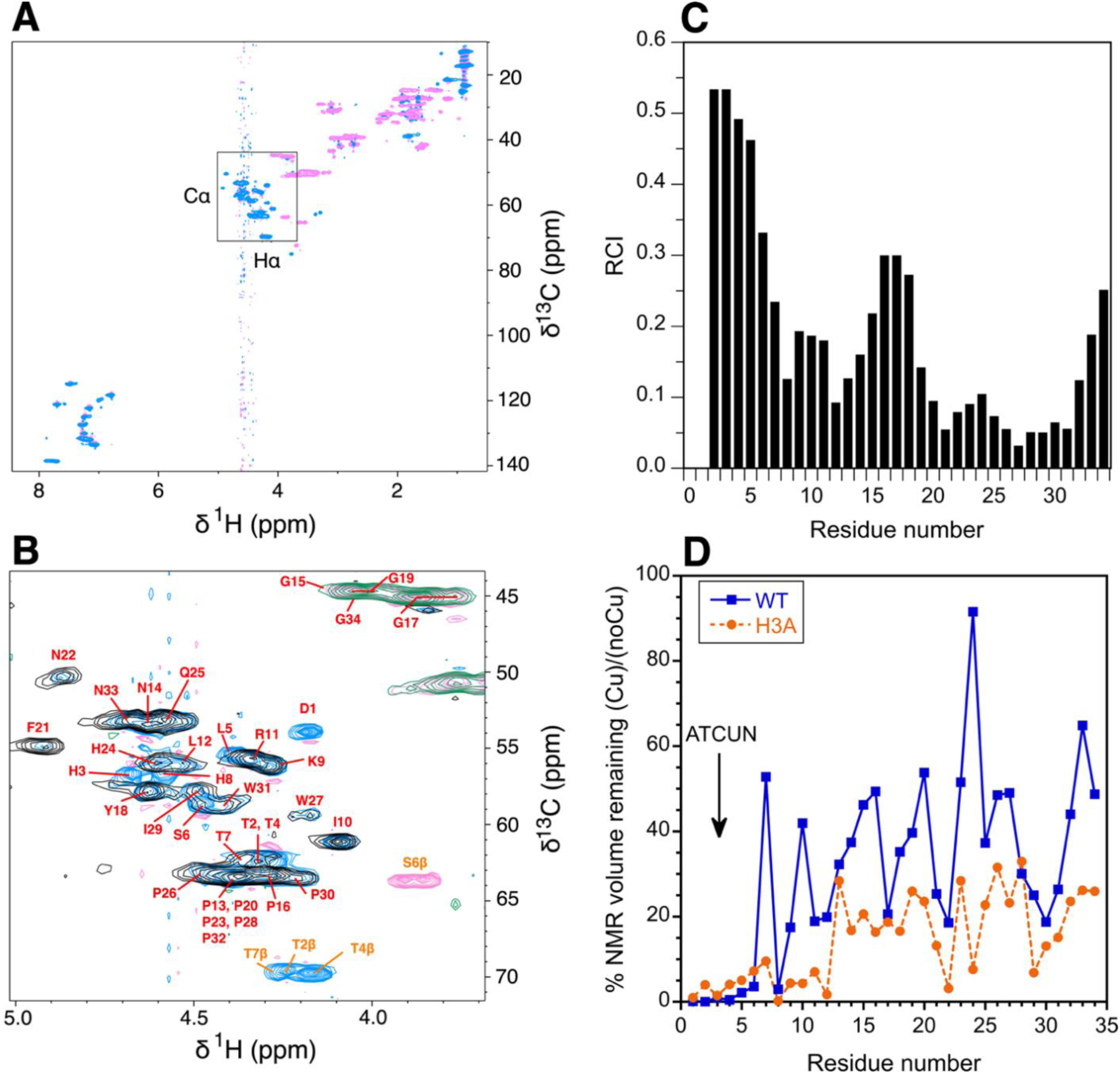
NMR analysis of Cu(II)-binding by WT abaecin-2. (**A**) A multiplicity-edited ^1^H-^13^C HSQC spectrum recorded at natural isotopic abundance for WT abaecin-2 from *A. texana*. In the multiplicity-edited spectrum, CH (aromatic and methine) and CH_3_ (methyl) groups are positive (blue contours), whereas CH_2_ groups (methylene) are negative (pink contours). The box shows the fingerprint region where each amino acid gives a Hα-Cα correlation used to analyze the paramagnetic Cu(II)-induced broadening on a per-residue basis. (**B**) Superposition of Hα-Cα regions for 1 mM WT abaecin-2 (blue and pink contours) and for the same peptide in the presence of 1.2 mM Cu(II) (black and green contours). Assignments for the CH groups of the WT abaecin-2 are labeled red, and those for the CH_2_ groups of Ser and Thr residues are orange. The spectrum with Cu(II) is shown at twice the amplitude to emphasize broadening. Note the selective broadening of peaks from the first nine residues in the presence of Cu(II). (**C**) Random coil index (RCI) calculated from the NMR assignments of the WT abaecin-2^32,46^. The larger the RCI, the more disordered the residue. RCI values for amino acids in folded regions of proteins are typically below 0.05. (**D**) Percent NMR cross-peak volume persisting after the addition of Cu(II). Cross-peak volumes were calculated from the NMR data in panel B for WT (squares, solid line) and H3A (circles, dotted line) abaecin-2. Note that broadening for WT abaecin-2 occurs primarily for the first nine residues, but for the H3A mutant peptide, the effects are more widespread, suggesting more non-specific interactions with Cu(II) for the mutant.

Based on these NMR chemical shift data, we calculated the random coil index (RCI)^32,46^ values for WT abaecin-2 (Figure 7C), where larger RCI values indicate a greater dynamic flexibility of a residue. The RCI values for the H3A mutant are very close to those of WT, indicating similar levels of disorder between the two peptides (Figure S10). In folded globular proteins, RCI values tend to be below ∼0.05^32,46,47^. The region from residues 20-32 shows lowered RCI values (relative to the rest of the peptide) near 0.1, likely due to the six prolines in this region increasing the rigidity of the polypeptide chain. Nevertheless, overall, our RCI analysis supports our conclusion based on CD and AF3 simulations that WT and H3A abaecin-2 peptides are largely unstructured in solution, although we cannot rule out that they become structured when they interact with other proteins or membranes in their physiological environment.

Binding of the paramagnetic metal Cu(II) causes a substantial decrease in backbone Hα-Cα NMR cross-peak volumes (typically to 0-5% of the original volume) for the first nine residues of the WT abaecin-2 peptide, which contains the ATCUN motif (Figure 7D). The aromatic ring resonances of H24, well outside the ATCUN motif, have the sidechain NMR signals that are broadened beyond detection (Figure S8), while the backbone signals are less affected (Figure 7B). Thus, His24 may represent a weak affinity site for Cu(II) that is responsible for super-stoichiometric metal binding in the WT abaecin-2. In the H3A variant, Cu(II)-induced NMR broadening is more nonspecific (Figure 7D, orange), with most residues experiencing a 70-80% loss in cross-peak volume, although the effects were still strongest in the ATCUN region. In both the WT and H3A peptides, NMR signals from the C-terminal residues in the presence of Cu(II) were superimposable with those obtained in the absence of that metal (Figure S8), indicating that metal binding does not affect the structure of the abaecin-2 C-terminus.

### Antibacterial susceptibility assays

To assess whether abaecin-2 displays antibacterial activity, we determined the minimum inhibitory concentration (MIC) of abaecin-2 peptides from *A. texana* against the Gram-positive bacterial species *Bacillus subtilis* 0114 and the Gram-negative bacterial species *Escherichia coli* MG1655. MIC values of >128 μM were obtained in all cases (Table 1, Table S4), suggesting that neither WT abaecin-2 nor the H3A variant, by themselves, had inhibitory activity against these bacteria. For comparison, piscidin-1 and piscidin-3, two well-studied antimicrobial HDPs, have MIC values that range from 2 to 8 μM depending on the bacterial strain used^40^. Also, the antimicrobial cecropin A and magainin-2 had MIC values between 1 and 8 μM in our study (Table 1). The lack of antimicrobial activity by abaecin-2 might be due to its moderate interaction with LPS compared to antimicrobial peptides such as cecropin A, magainin-2, and polymyxin B (Figure S11) ^39,51,52^. However, strong interaction with LPS is not a requirement for antimicrobial activity, as observed in the case of streptomycin (Figure S11).

**Table 1.**
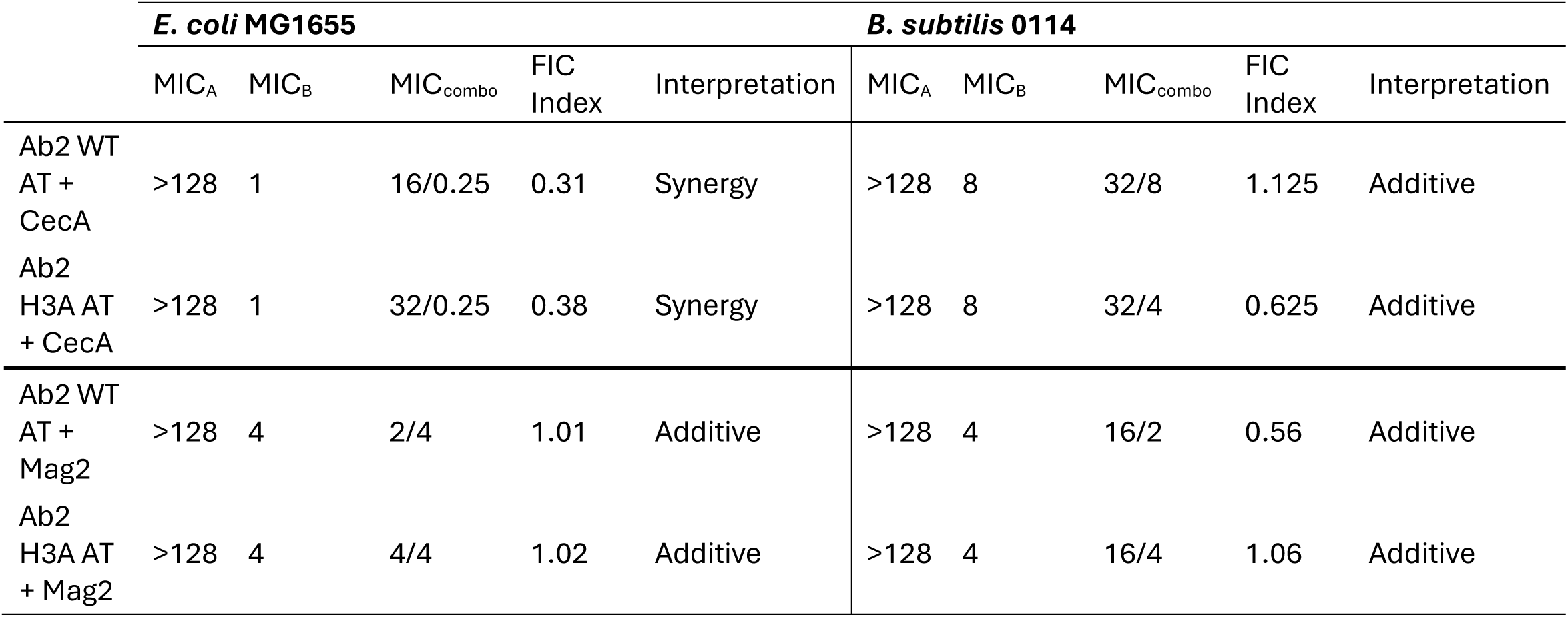
Antibacterial susceptibility assays. WT and H3A abaecin-2 (Ab2) peptides from *A. texana* were assayed, with and without the pore-forming peptides cecropin A (CecA) or magainin-2 (Mag2), against *E. coli* MG1655 or *B. subtilis* 0114. MIC_A_, MIC_B_, and MIC_combo_ indicate, respectively, the MIC when abaecin-2 alone, either cecropin A or magainin-2 alone, or a combination was assayed together in MHB.

Abaecin-2, like abaecin-1^15,16^, may require a pore-forming partner peptide for antibacterial activity. We therefore measured the fractional inhibitory concentration (FIC)^36^ of *A. texana* abaecin-2 in combination with cecropin A or magainin-2, two well-characterized α-helical, hydrophobic, pore-forming peptides from the moth *Hylaphora cecropia* and the African Clawed Frog *Xenopus laevis*, respectively.^48,49^ As hypothesized, cecropin A synergized with both WT and H3A abaecin-2 peptides to inhibit *E. coli* growth, but magainin-2 only interacted with abaecin-2 additively (Table 1, Table S4). These results are similar to those obtained using abaecin-1, in which only cecropin A showed synergistic behavior^16^. In the case of *B. subtilis* 0114, both cecropin A and magainin-2 had additive effects with abaecin-2 (Table 1). The two pore-forming peptides have physicochemical differences in their electrostatic charges and Boman index (Table 2)^50^, although it is not clear how these characteristics may influence synergy. More importantly, no major differences were observed in the FIC values for WT or H3A abaecin-2, indicating that the potentiating effect of the pore-forming peptides does not depend on an intact ATCUN motif.

**Table 2.** Biochemical properties of peptides in this study. The ATCUN motif is indicated in bold, and a predicted DnaK binding motif is underlined. The (G) indicates the addition of a glycine to the C-terminus to simplify synthesis.

| Peptide | Sequence | Net Charge<br>(pH = 7.0) | Hydrophobicity<br>% | Boman Index |
| --- | --- | --- | --- | --- |
| Abaecin-2, *A. texana* | **D**<u>**T**H**T**L**S**T**H**K**I**R**L**P**N**G**P**G**Y**G**P**F**N**H**Q**P**W**P**I**P**W**P**N**</u>(G)-NH<sub>2</sub> | 2.3 | 24 | - |
| Abaecin-2, *A. colombica* | **D**<u>**T**H**T**F**S**T**H**K**I**R**L**P**N**G**P**G**Y**G**P**F**N**H**Q**P**W**P**I**P**W**P**N**</u>(G)-NH<sub>2</sub> | 2.27 | 24 | - |
| Cecropin A | KWKLFFKKIEKVGQNIRDGIIKAGPAVAVVGQATQIAK-NH<sub>2</sub> | 7 | 45 | 0.84 |
| Magainin-2 | GIGKFLHSAKKFGKAFVGEIMNS-NH<sub>2</sub> | 4.1 | 43 | 0.41 |

We also investigated if the pore-forming activity of cecropin A allows abacein-2 to reach and inhibit an intracellular target. The sequence of *A. texana* abaecin-2 (Table 2) contains a non-canonical DnaK binding motif that is similar to the motif in bumblebee abaecin-1 (WPYPLPN)^15^. To examine if abaecin-2 targets DnaK, we assessed the MIC for this peptide against both WT *E. coli* and a Δ*dnaK* mutant strain, which lacks the DnaK protein. If DnaK was indeed the target, abaecin-2 would be expected to have a higher MIC (indicating loss of activity) and higher FIC (indicating loss of synergy) with cecropin A against the Δ*dnaK* strain compared to the WT strain. We did not observe a difference (Table 1, Table 3). This suggests that abaecin-2 has an unknown intracellular target that is distinct from that of abaecin-1.

**Table 3.** Susceptibility of *E. coli* JW 0013-4 (Δ*dnaK*) to *A. texana* abaecin-2 variants in combination with the pore-forming peptide cecropin A. All assays were conducted in M9 medium.

| | *E. coli* JW 0013-4 ($\Delta dnaK$) | | | | |
| --- | --- | --- | --- | --- | --- |
|  | MIC<sub>A</sub> | MIC<sub>B</sub> | MIC<sub>combo</sub> | FIC Index | Interpretation |
| Abaecin-2 WT AT + CecA | >128 | 1 | 16/0.25 | 0.31 | Synergy |
| Abaecin-2 H3A AT + CecA | >128 | 1 | 16/0.25 | 0.31 | Synergy |

### Abaecin-2 rescues copper-sensitive model bacteria

Our data indicate that abaecin-2 binds Cu(II) ions. However, the ATCUN motif is not essential for synergy with cecropin A. ATCUN-containing HDPs typically use the ATCUN-Cu(II) complex to produce reactive oxygen species (ROS) to damage their targets^2,40,41,53^, while other ATCUN motifs inhibit the formation of ROS upon formation of the Cu(II) complex. For example, ATCUN motifs containing a Glu or Asp residue in position 1 produce ROS with at least an order of magnitude less efficiency than other ATCUN motifs that contain hydrophobic residues at this position^54^. Because the ATCUN motif in mymicine ants is either a DTH or DIH motif, we hypothesized that the ATCUN motif in these peptides could instead protect symbionts against copper toxicity, possibly to promote colony and fungus garden health.

To investigate abaecin-2’s potential protective role in preventing copper toxicity in symbionts sensitive to copper, we studied the effect of abaecin-2 supplementation on the growth of a model bacterial strain, a copper-sensitive *E. coli* Δ*copA* mutant that lacks the primary copper efflux pump, grown in the presence of different levels of copper. The WT abaecin-2 alone does not inhibit *E. coli* Δ*copA* (MIC >128 μM; Table S4). Our initial experiments used Mueller Hinton Broth (MHB). Minimal M9 medium was used to assess the protective effects of abaecin-2 during copper toxicity to avoid background interference from other copper-binding molecules in MHB. Copper inhibited both WT and Δ*copA E. coli* strains in a dose-dependent manner (Figure 8). As expected, the Δ*copA* strain was more sensitive (Figure 8) to copper stress, compared to exposure of these strains to Cu(II) alone. The presence of 10 μM WT abaecin-2 rescued both strains, consistent with a protective effect. Interestingly, the H3A mutant (also 10 μM) was also protective, although only to a lower concentration of Cu(II) (up to 1 μM; Figure S12).

**Figure 8.**
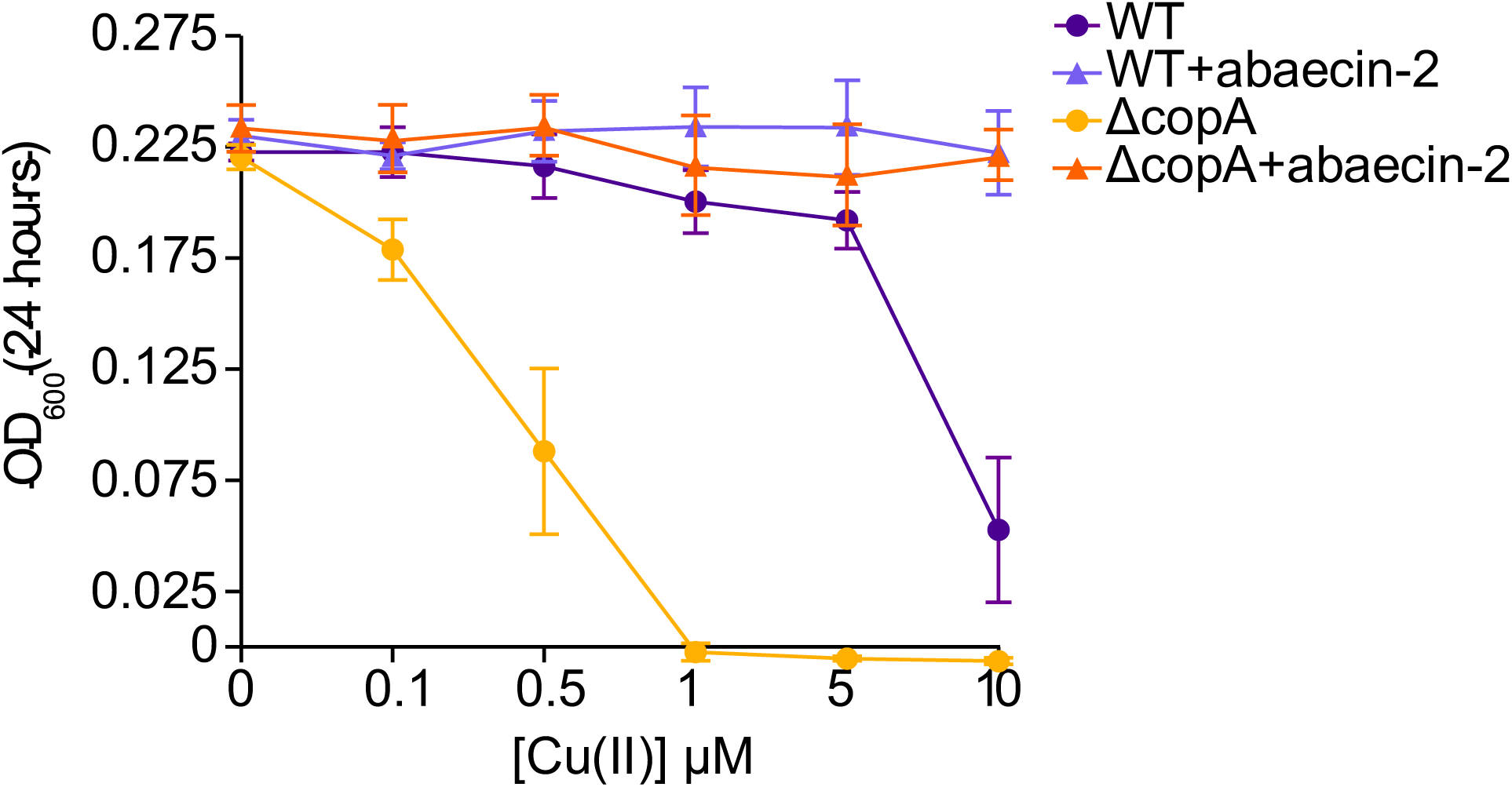
Abaecin-2 rescues *E. coli* from copper toxicity. Data points show growth of either WT or the Δ*copA* mutant strain of *E. coli*, represented by the Optical Density at 600 nm after 24 h incubation at 37 °C, in the presence of Cu(II), with or without 10 μM WT *A. texana* abaecin-2.

## Conclusion

In this work, we described the structure and potential mechanisms of action of an uncharacterized HDP, abaecin-2, from several fungus-growing ant species. Although abaecin-2 first evolved before the emergence of ants, it only later evolved a copper-binding ATCUN motif that, along with additional His residues, is conserved in fungus-growing ants. Abaecin-2 from *A. texana* and *A. colombica* binds Cu(II) with a high affinity at the ATCUN motif, with weaker binding at the other conserved His residues. The growth of model Gram-positive and Gram-negative bacteria are not inhibited by abaecin-2 alone. However, the presence of the membrane-disrupting peptide cecropin A leads to synergistic antibacterial activity against *E. coli*. Our results indicate that the ATCUN motif is not involved in this synergistic behavior. In the absence of additional HDPs, binding of Cu(II) by abaecin-2 can also prevent copper toxicity in a model copper-sensitive mutant *E. coli* strain, raising the possibility that it performs a similar function in wild fungus gardens, which are predominantly colonized by related bacteria in the family *Enterobacteriaceae*^55,56^. The dual activities of abaecin-2 towards the model Gram-negative bacterial species *E. coli*—antibacterial activity potentiated by additional peptides or rescue by binding excess copper—indicate that abaecin-2 may regulate microbial symbionts within fungus-growing ant colonies. We note that this current study leveraged the experimental tractability of model partner peptides and target microbes, and that these may differ somewhat from those present in vivo. Having established foundational properties of abaecin-2, our future work will focus on these more complex in vivo interactions, e.g., including the full diversity of beneficial and pathogenic bacteria and fungi in the fungus-growing ant symbiosis, and involving the pore-forming HDPs encoded by fungus-growing ant genomes, e.g., the structurally complex HDP hymenoptaecin^13^. Doing so will reveal the natural conditions in which abaecin-2 synergistically increases antimicrobial activity or protects various microbial symbionts from copper toxicity.

## Acknowledgements

Funding for this study was provided via an NSF Graduate Research Fellowship Program award to C.D. under Grant No. DGE-17474753, a UConn Convergence Awards for Research in Interdisciplinary Centers grant and a Microbiome Seed Grant to J.K. and A.A.-B, and a Durham University Seedcorn Grant to K.D. We thank Emily Green and Helen Edwards for their assistance with ant collections, and Samantha Firth and Emma Tarrant for assistance with ICP-MS measurements at Durham. NMR experiments were done at the Gregory P. Mullen NMR Structural Biology Facility of UConn Health, which is a member of the NSF Network for Advanced NMR (grants 1946970 and 2529058). Their 800 MHz and 700 MHz instruments were funded by NIH grants S10RR023041 and S10OD034297, respectively.

