## Supplementary Figures and Tables for "Ant abaecin-2 is a context-dependent copper-binding effector that can be either inhibitory or protective"

**Table of Contents:**

**Figure S1:** Maximum likelihood tree of ant abaecin-2 protein sequences.
**Figure S2:** Maximum likelihood tree of ant abaecin-2 nucleotide sequences.
**Figure S3.** Mass spectrometry of abaecin-2 peptides to determine their copper-binding ability.
**Figure S4:** Copper recovery from *A. colombica* abaecin-2 after desalting.
**Figure S5:** Cu(II) binding affinity of *A. colombica* abaecin-2.
**Figure S6:** 1D NMR spectra of abaecin-2 WT and H3 illustrating Cu(II) induced line broadening.
**Figure S7:** NMR broadening of WT-abaecin-2 TOCSY cross-peaks with Cu(II) affects primarily the first nine residues in the sequence.
**Figure S8:** Superposition of NMR spectra illustrating differences due to the H3A mutation between WT-abaecin-2 and H3A-abaecin-2.
**Figure S9:** Multiplicity-edited $^{13}$C-HSQC NMR spectra illustrating signal broadening due to Cu(II) in WT-abaecin-2 and H3A-abaecin-2.
**Figure S10.** Comparison of random coil index (RCI) values for abaecin-2 WT and the H3A mutant.
**Figure S11.** Interaction of *A. texana* abaecin-2 with LPS
**Figure S12.** Differences in the rescuing effect of *A. texana* WT and H3A abaecin-2 peptides.
**Table S1:** Abaecin-2 sequences used for BLASTp searches.
**Table S2:** Hymenopteran abaecin sequence data returned by BLAST searches.
**Table S3:** ICP analysis sample metadata.
**Table S4.** Susceptibility of *E. coli* JW 0473-3 (Δ*copA*) to *A. texana* and *A. colombica* abaecin-2 variants in combination with the pore-forming peptide cecropin A.
**Supplemental code S1.** Sample DynaFit scripts used for Cu(II) binding affinity determination of abaecin-2 peptides.

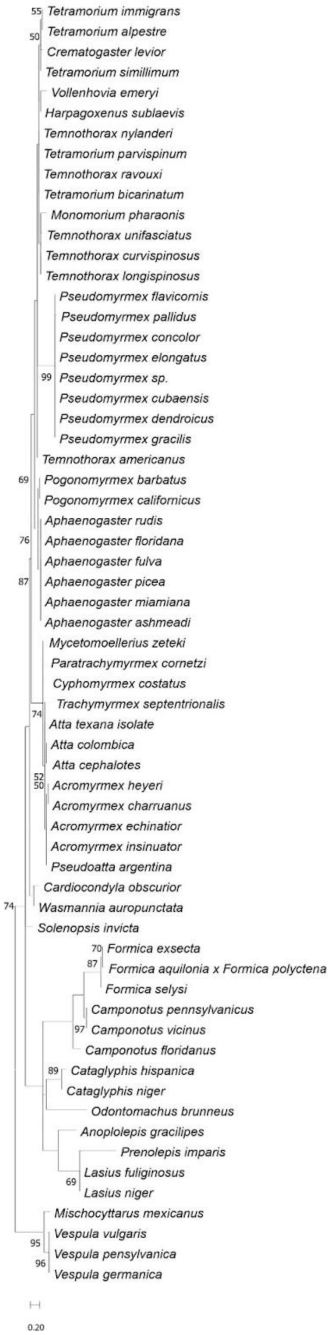


**Figure S1. Maximum likelihood tree of ant abaecin-2 protein sequences.** Protein phylogeny generated from abaecin-2 peptide sequences using the MEGA JTT+G substitution model with 500 bootstrap replicates. Abaecin-2 sequences from 4 wasp species (*Mischocyttarus mexicanus, Vespula vulgaris, V. pensylvanica,* and *V. germanica*) were used as an outgroup to root the tree. Local bootstrap values > 50 are shown. Scale bar corresponds to the number of amino acid substitutions per site.

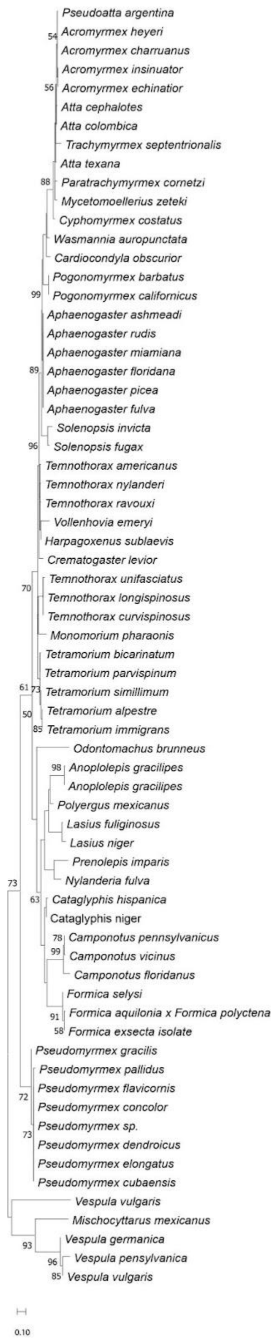


**Figure S2. Maximum likelihood tree of ant abaecin-2 nucleotide sequences.** Nucleotide phylogeny generated from abaecin-2 coding nucleotide sequences using the MEGA K2 substitution model with 500 bootstrap replicates. Abaecin-2 nucleotide sequences from 4 wasp species (*Mischocyttarus mexicanus, Vespula vulgaris, V. pensylvanica,* and *V. germanica*) were used as an outgroup to root the tree. Local bootstrap values > 50 are shown. Scale bar corresponds to the number of nucleotide substitutions per site.

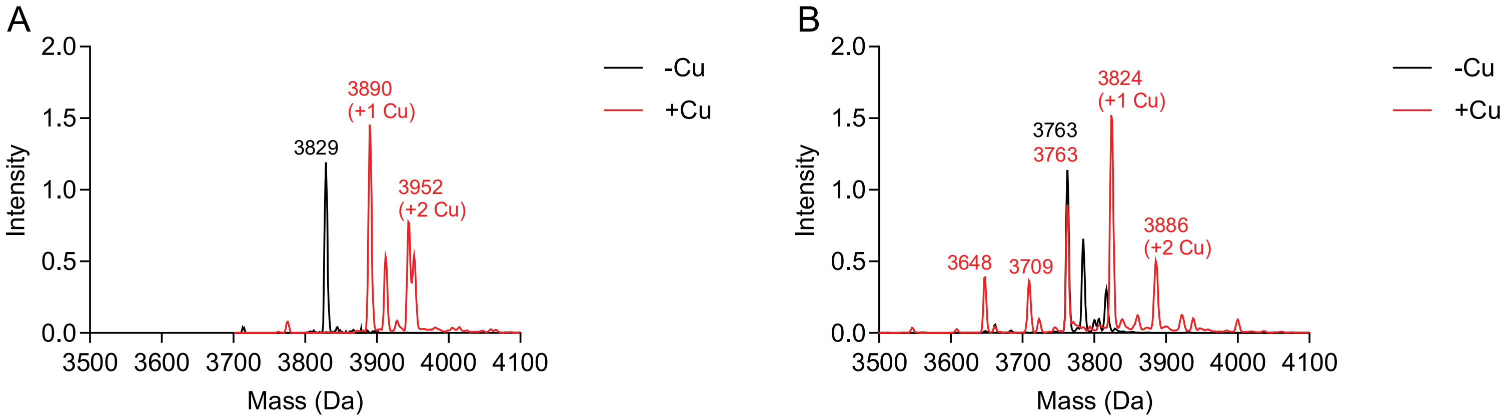


**Figure S3. Mass spectrometry of abaecin-2 peptides to determine their copper-binding ability.** (A) Wildtype (WT) *A. texana* abaecin-2 incubated in the presence or absence of 5 molar equivalents of Cu(II). (B) H3A *A. texana* abaecin-2 incubated in the presence or absence of 5 molar equivalents of Cu(II). Both peptides can bind up to two Cu(II) ions, but binding is weaker in the H3A mutant.

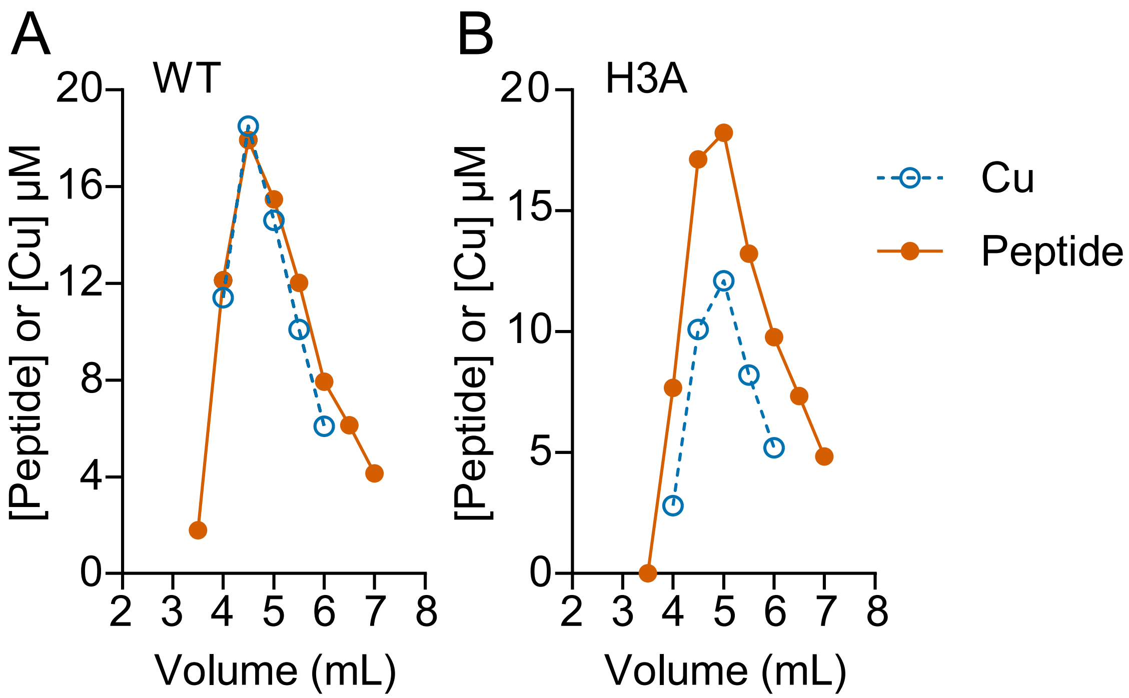


**Figure S4. Copper recovery from *A. colombica* abaecin-2 after desalting.** Elution of WT (A) and H3A (B) abaecin-2 peptides from PD-10 desalting column. Data points show the concentrations of peptide (filled circles, ●) and Cu(II) ions (open circles, ○) in each fraction. Data are from one representative of three independent experiments.

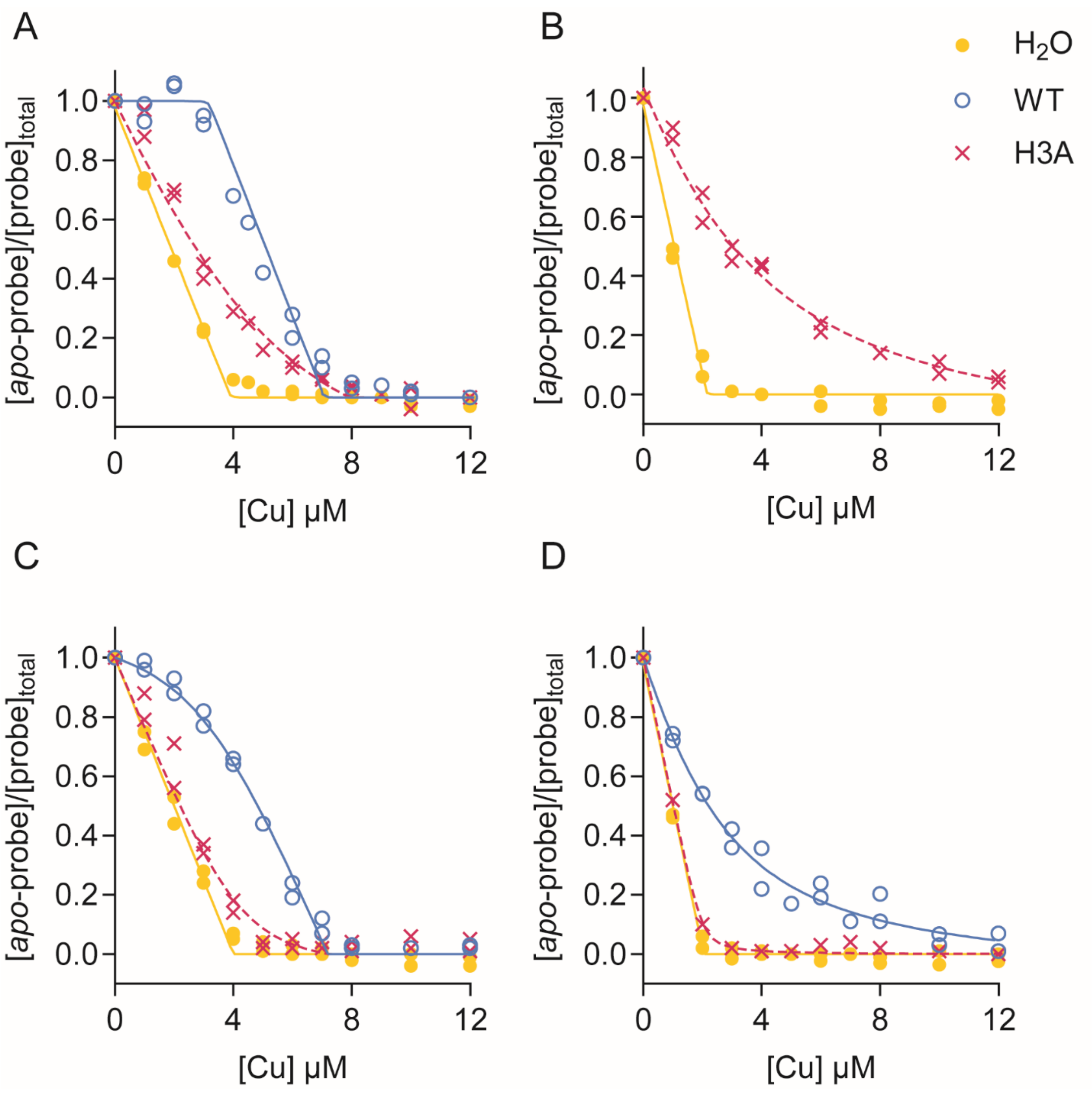


**Figure S5. Cu(II) binding affinity of *A. colombica* abaecin-2.** The WT (○, solid lines) or H3A (x, dashed lines) peptides were competed against the fluorescence probe DP2, DP3, or DP4. Data points (n = 2) show the resulting competition curves when Cu(II) was added to mixtures of (A) 4 µM DP2 and 3.2 µM peptide, (B) 2 µM DP2 and 12.8 µM peptide, (C) 4 µM DP3 and 3.2 µM peptide, or (D) 2 µM DP4 and peptide. Control curves without any peptide are also shown (●, solid lines).

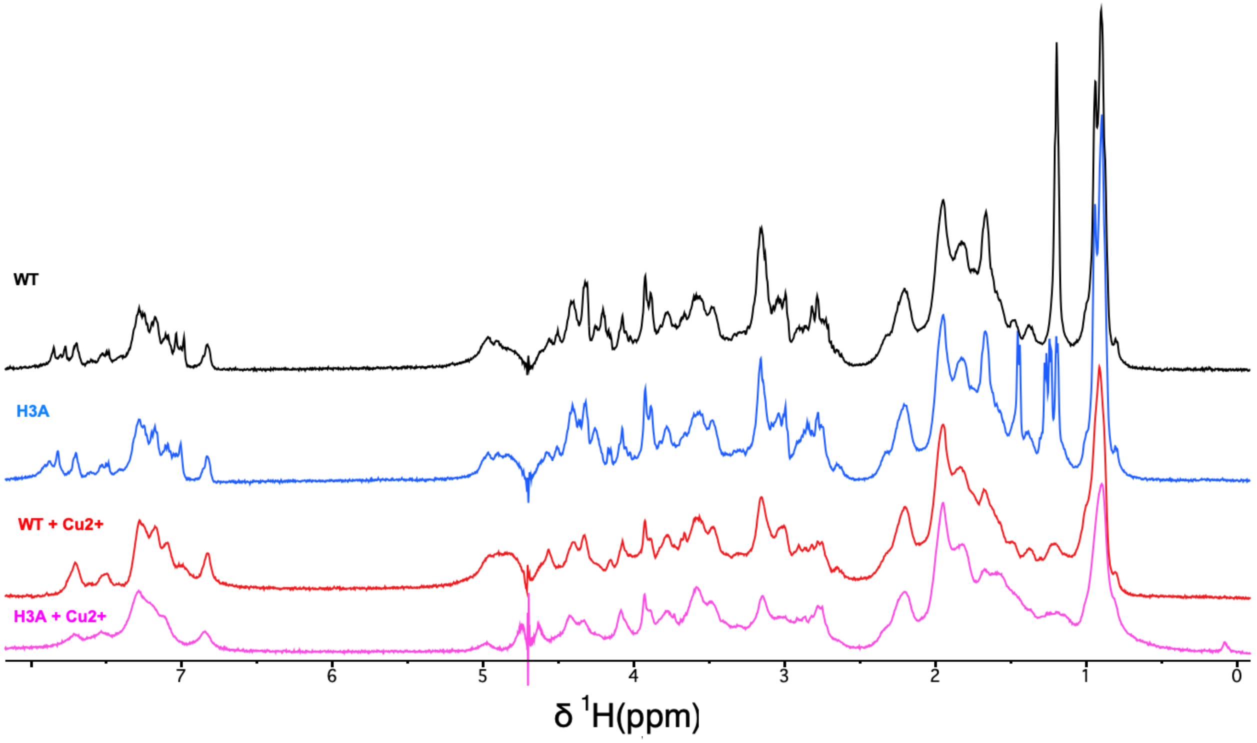


**Figure S6: 1D NMR spectra of abaecin-2 WT and H3A illustrating Cu(II) induced line broadening.** WT and H3A concentrations were 0.5 mM in the presence or absence of 0.6 mM $CuCl_2$ (1.2 molar equivalents to abaecin-2). Data were collected at 500 MHz and a temperature of 32 °C for samples in $D_2O$, with a pD of 6.6 ± 0.2.

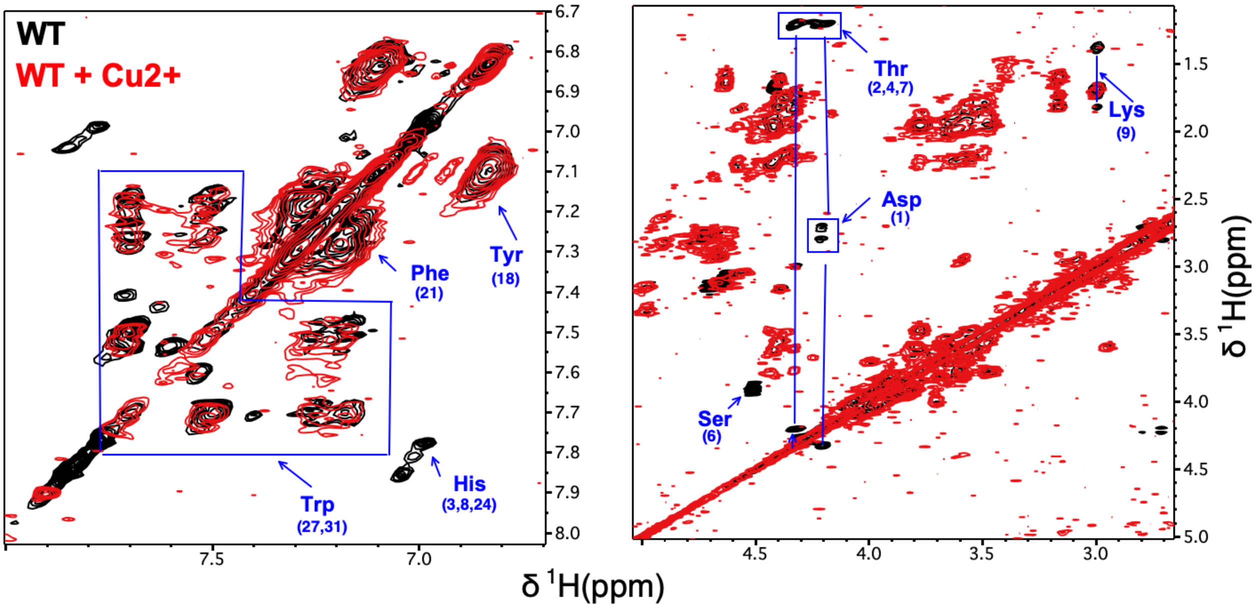


**Figure S7: NMR broadening of WT-abaecin-2 TOCSY cross-peaks with Cu(II) affects primarily the first nine residues in the sequence.** Superposition of 70 ms mixing time 2D TOCSY spectra for WT-abaecin-2 in the presence (red) or absence (black) of 1.2 molar equivalents of $CuCl_2$. Left and right panels show the aromatic and aliphatic portions of the spectra. Most of the spin systems affected by Cu(II) are in the first nine residues (sequence numbers are given in parentheses), whereas the other cross-peaks are largely unaffected. The exceptions are the Hε1-Hδ2 cross-peaks of H24, and two sidechain cross-peaks from either W27 or W31 that are also absent in the presence of Cu(II). TOCSY spectra were recorded at 500 MHz on a 0.5 mM WT-abaecin-2 sample at pD 6.6.

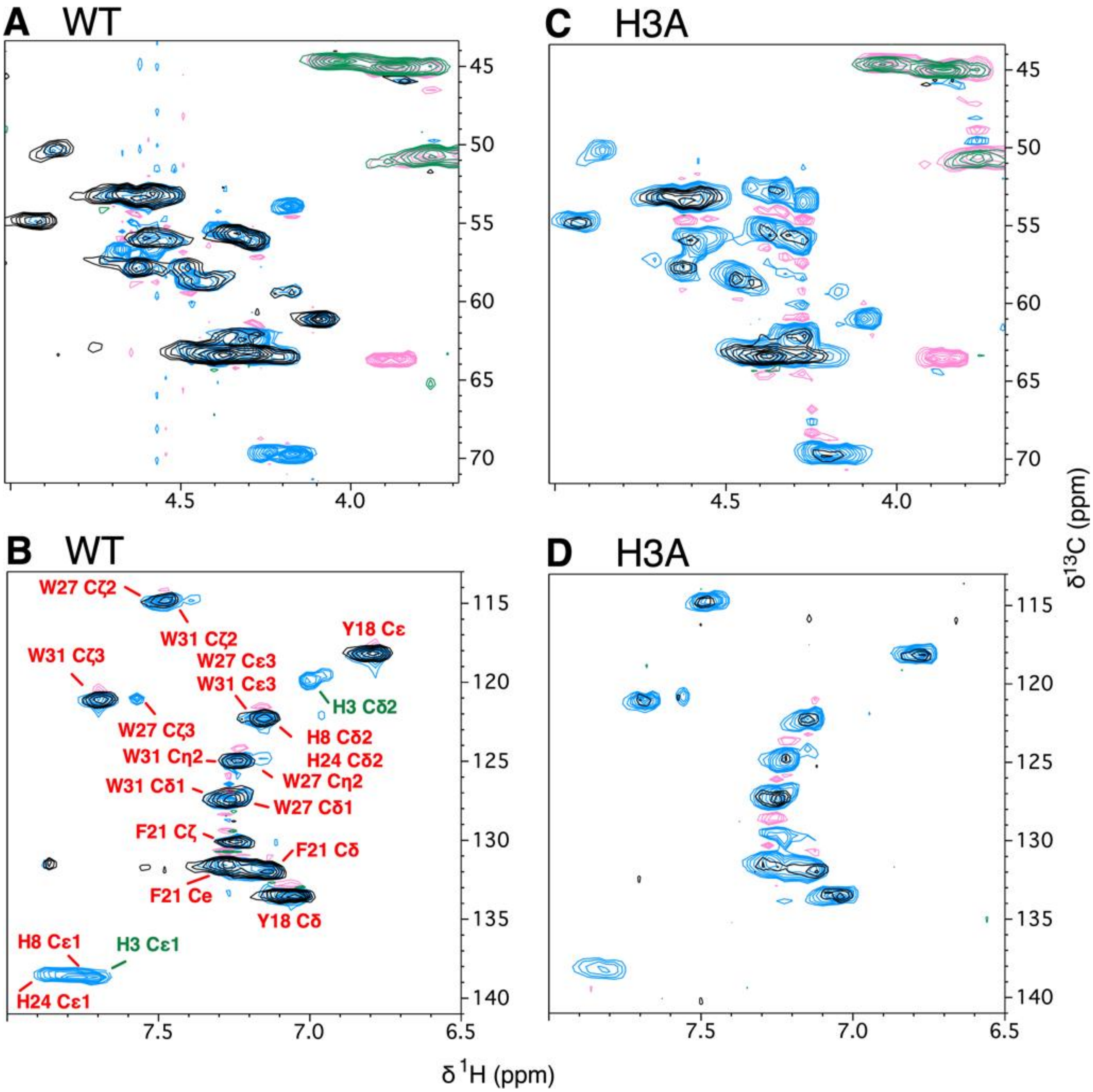


**Figure S8: Multiplicity-edited $^{13}$C-HSQC NMR spectra illustrating NMR signal broadening due to Cu(II) in abaecein-2 WT and the H3A mutant.** In the multiplicity-edited $^{13}$C-HSQC experiments, CH and $CH_3$ groups have positive contours while $CH_2$ groups have negative contours. The Hα-Cα fingerprint regions and aromatic regions of the spectrum respectively are shown for WT (**A**, **B**) and H3A (**C**, **D**). For the spectra in the absence of Cu(II), contour levels are shown in blue (positive) and pink (negative). The spectra in the presence of Cu(II) are shown with black (positive) and green (negative) contours. For all panels, the spectra in the presence of Cu(II) are shown at twice the amplitude to emphasize the line broadening. The assignments for the Hα-Cα region are given in Figure 7B of the main text. The assignments for the aromatic region are given in panel B of this figure. The multiplicity-edited $^{13}$C spectra were collected at 800 MHz at a temperature of 10 °C. WT and H3A concentrations were 1 mM in the presence or absence of 1.2 mM $CuCl_2$ (1.2 molar equivalents to abaecin-2). Note that Cu(II)-induced NMR broadening is more widespread in H3A and more specific in WT.

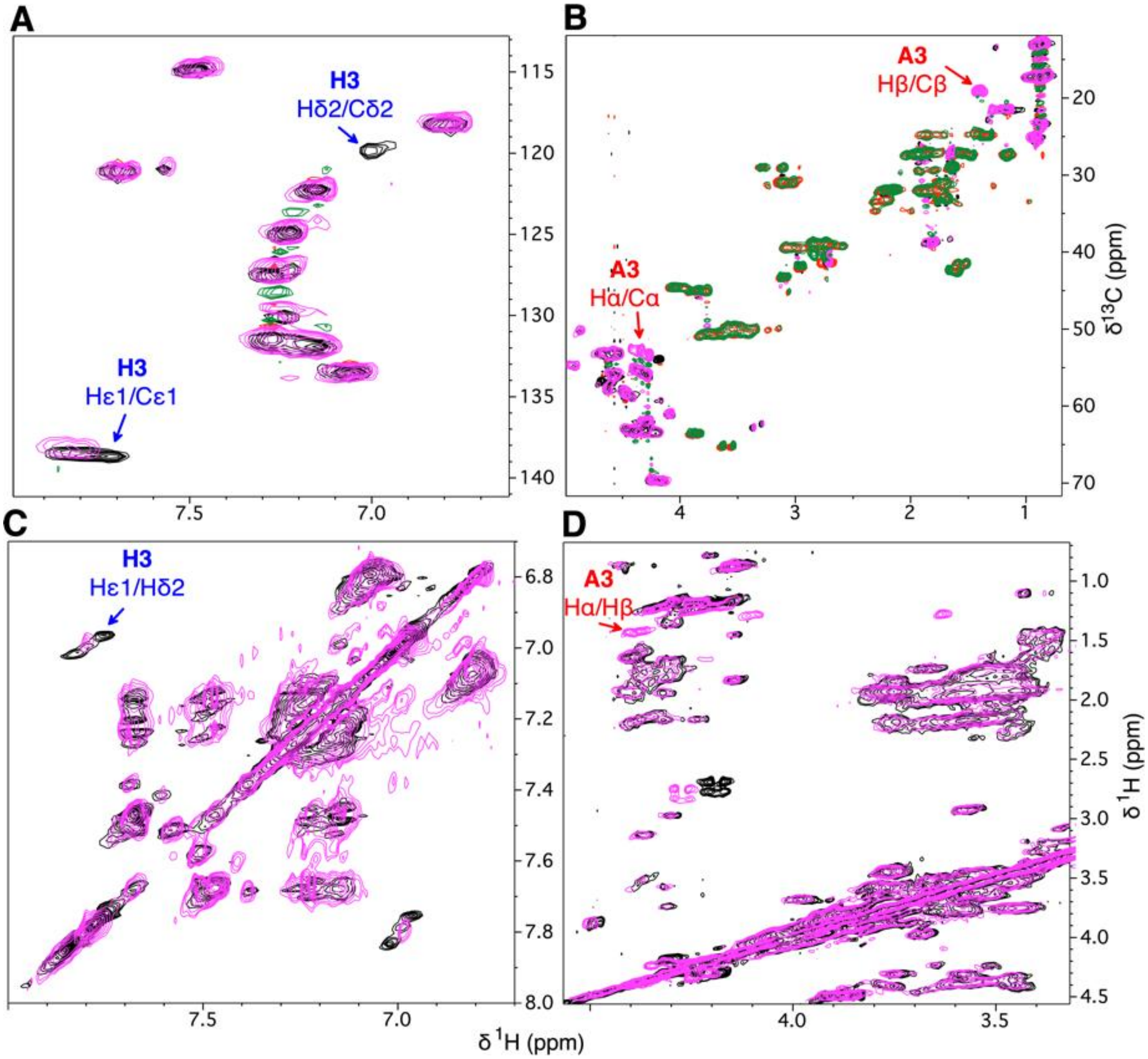


**Figure S9: Superposition of NMR spectra illustrating differences due to the H3A mutation in abaecin-2.** Aromatic (**A**) and aliphatic (**B**) regions of the superposed multiplicity-edited $^{13}$C-HSQC spectra of WT (black for positive contours from CH and $CH_3$ groups, red for negative contours from $CH_2$ groups) and H3A (pink for positive contours and green for negative contours). The extra black contours due to the Hε1/Cε1 and Hδ2/Cδ2 cross-peaks of H3 in WT compared to H3A are labeled blue in (A). Similarly, the extra pink contours due to the Hα/Cα and Hβ/Cβ cross-peaks of A3 in H3A compared to WT are labeled red in (B). The bottom two panels show superposed aromatic (**C**) and aliphatic (**D**) regions from the 2D TOCSY spectra for WT (black) and H3A (pink). The differences between the two spectra due to the presence of H3 in WT and A3 in H3A, are labeled blue and red, respectively. The multiplicity-edited $^{13}$C-HSQC spectra were collected at 800 MHz using 1 mM samples at a temperature of 10 °C. The TOCSY data were collected at 500 MHz using 0.5 mM samples at a temperature of 32 °C.

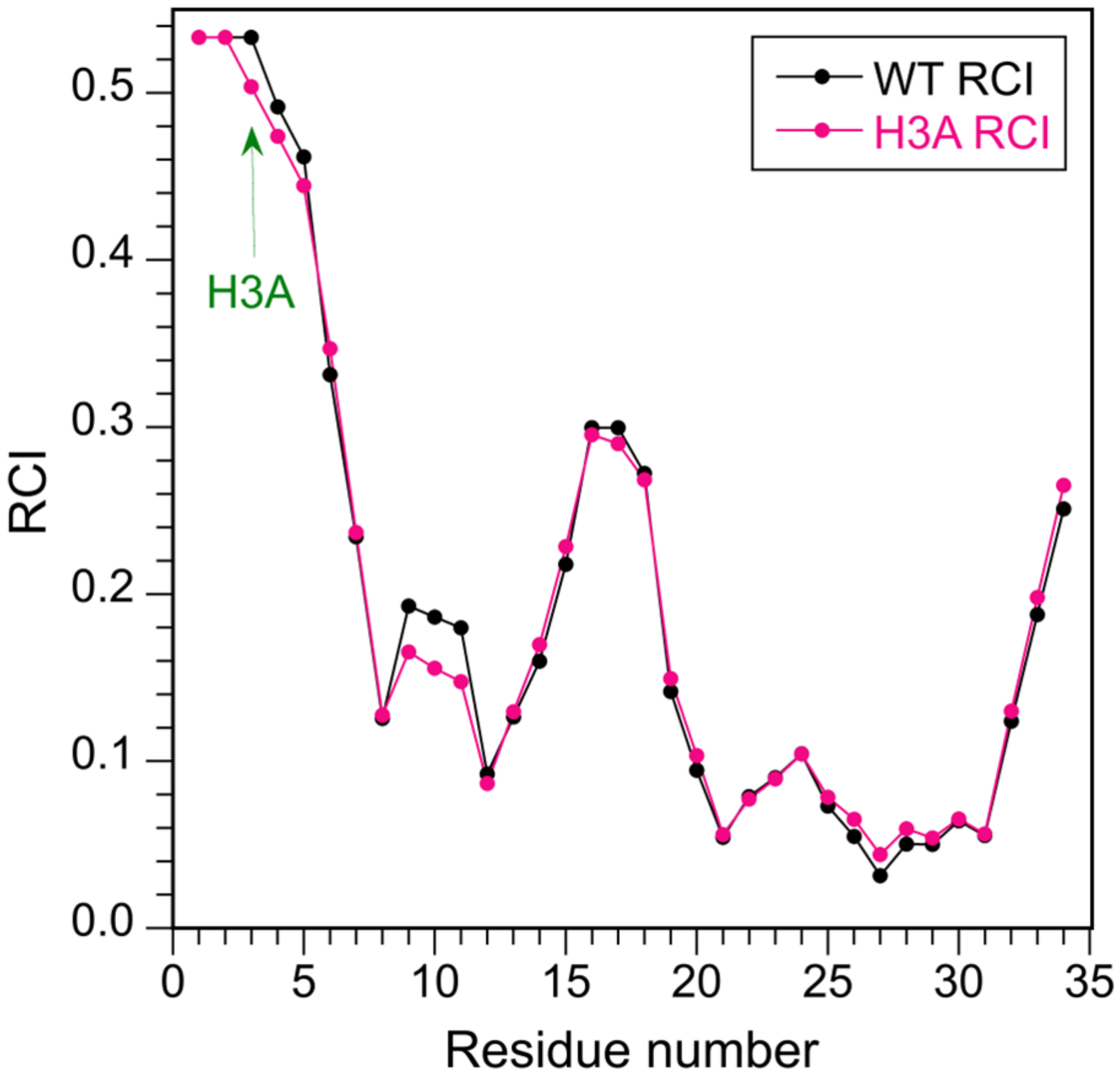


**Figure S10. Comparison of random coil index (RCI) values for abaecin-2 WT and the H3A mutant.** The site of the H3A mutation is denoted with a green arrow in the plot. The RCI sequence profiles are very close for the WT and mutant, consistent with their $^{1}H$-$^{13}C$ HSQC and TOCSY spectra being superposable (Fig. S9), and indicate similar levels of disorder.

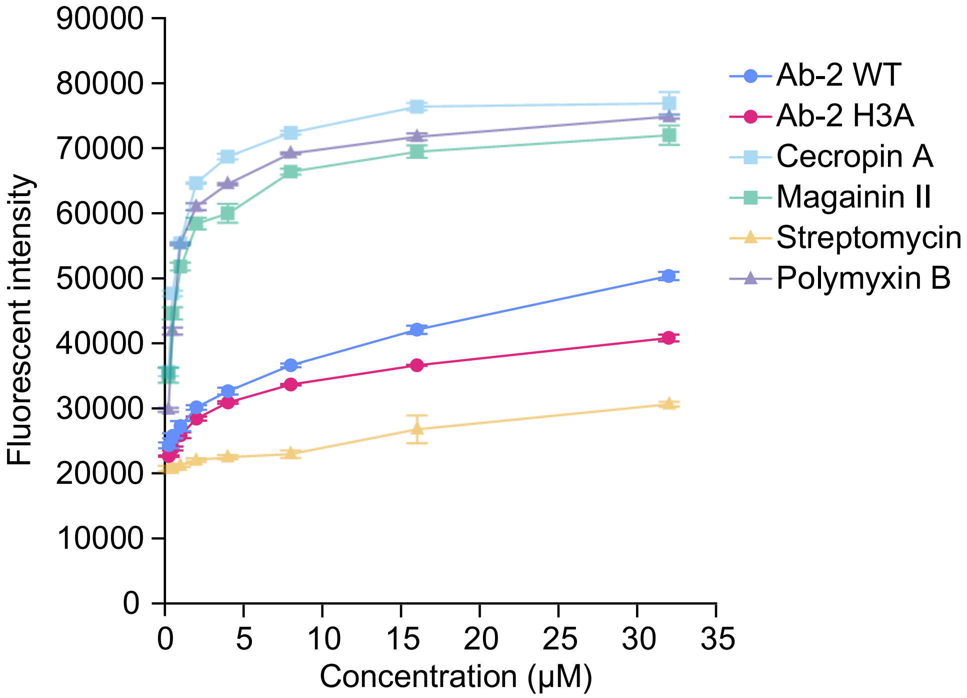


**Figure S11. LPS binding by abaecin-2 peptides, pore-forming peptides, and antibiotic controls.** Data points show the average (± SE from 2 replicates) fluorescent intensity observed during BODIPY-TR-cadaverine displacement assays. Both WT and H3A abaecin-2 peptides interact only moderately with LPS, unlike magainin-2, cecropin A, and polymyxin B.

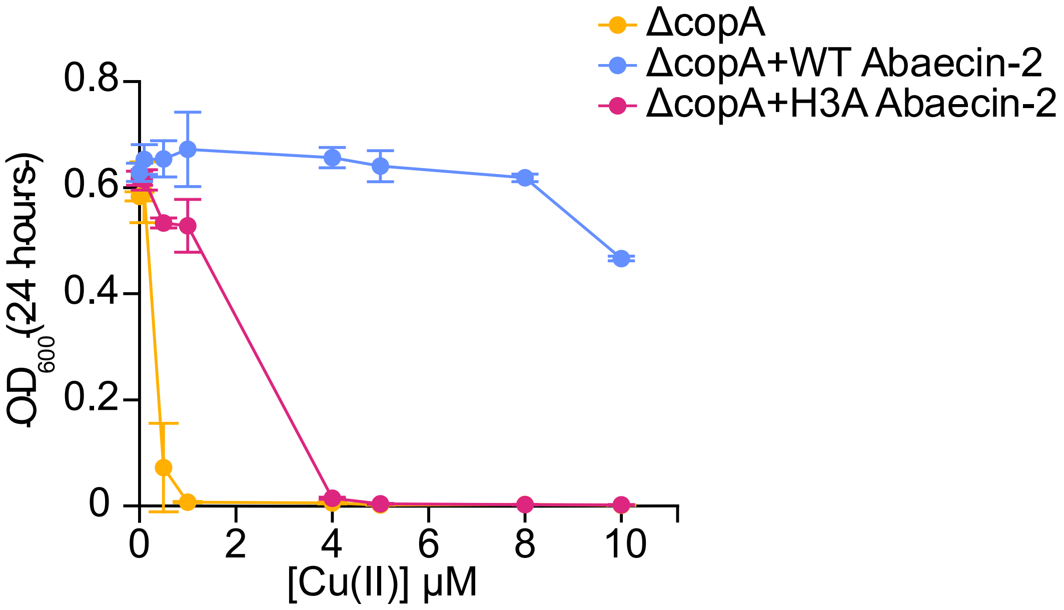


**Figure S12. Differences in the rescuing effect of *A. texana* WT and H3A abaecin-2 peptides.** Data points show growth of either the Δ*copA* mutant strain of *E. coli*, represented by the Optical Density at 600 nm after 24 h incubation at 37 °C, in the presence of Cu(II), with or without 10 μM of WT or H3A *A. texana* abaecin-2.

**Table S1. Abaecin-2 sequences used for BLASTp searches.**

| Species | Abaecin-2 sequence |
|---|---|
| *A. colombica* | DTHTFSTHKIRLPNGPGYGPFNPHQPWPIPWPNG |
| *A. texana* | DTHTLSTHKIRLPNGPGYGPFNPHQPWPIPWPNG |
| *Mycetomoellerius zeteki* | MRFTMYIVMLLVAICALFADTHTLLTHRIRLPNGPGYGPFNPHQPWPIPWPNNG |
| *Acromyrmex echinatior* | MRLTMYIVMLLITICALFADIHTLSTHKIRLPNGPGYGPFNPHQPWPIPWPNNG |
| *Wasmannia auropunctata* | MKFAMYMVMLLVAICALFADTHALPKHRFRLPSGPGYGPFNPQQPWPVPWPNNG |
| *Pseudomyrmex gracilis* | FKNNTHIVNMKFAIYAVMLLVAICALFANTQALPGRRSSLPCGPGYGPFNPRKSWPIPLPKHG |
| *Pogonomyrmex barbatus* | MRFSFPITTSFISLQSNIFNMRFAAYIVTLLVAICALFADTHALPKHRLRLPGGGPGYGPFNPRLPWPIPLPNRDH |
| *Cyphomyrmex costatus* | FYFPVTIFFDQNNIFNMRLRFTMYIVMLLVAICALFVDTHTLSTHRIRLPSGPGYGPFNPHQPWLISWPNQ |
| *Paratrachymyrmex cornetzi* | MRFTMYIVMLLIATCALFVDTHTLSTHRIRLPNGPGYGPFNPHQSWPIPLPNNG |

### Table S2. Hymenopteran abaecin sequence data returned by BLAST searches.

| Name | Organism | Accession number | Contig position | Abaecin type | Search method | Annotated gene model present?* |
|---|---|---|---|---|---|---|
| *Pogonomyrmex barbatus* | Ant | XP 025074443.1 | 1-49 | Abaecin-1 | PSI-BLAST Search Result | No |
| *Acromyrmex echinatior* | Ant | EGI68768.1 | 1-51 | Abaecin-1 | PSI-BLAST Search Result | No |
| *Atta colombica* | Ant | KYM77085.1 | 1-51 | Abaecin-1 | PSI-BLAST Search Result | No |
| *Cyphomyrmex costatus* | Ant | KYM99866.1 | 1-58 | Abaecin-1 | PSI-BLAST Search Result | No |
| *Harpegnathos saltator* | Ant | EFN88303.1 | 49-97 | Abaecin-1 | Translated BLASTn Search Result | No |
| *Mycetomoellerius zeteki* | Ant | KYQ52474.1 | 1-51 | Abaecin-1 | PSI-BLAST Search Result | No |
| *Ooceraea biroi* | Ant | EZA47299.1 | 1-52 | Abaecin-1 | Translated BLASTn Search Result | No |
| *Paratrachymyrmex cornetzi* | Ant | KYN17873.1 | 1-51 | Abaecin-1 | PSI-BLAST Search Result | No |
| *Solenopsis invicta* | Ant | AFJ94682.1 | 1-51 | Abaecin-1 | Translated BLASTn Search Result | No |
| *Temnothorax curvispinosus* | Ant | XP 024869487.1 | 1-52 | Abaecin-1 | Translated BLASTn Search Result | No |
| *Apis cerana japonica* | Bee | BAI58048.1 | 1-53 | Abaecin-1 | Translated BLASTn Search Result | No |
| *Apis cerana* | Bee | XP 016919244.1 | 1-53 | Abaecin-1 | Translated BLASTn Search Result | No |
| *Apis cerana cerana* | Bee | ACH90419.1 | 1-53 | Abaecin-1 | Translated BLASTn Search Result | No |
| *Apis florea* | Bee | XP 003699027.2 | 21-78 | Abaecin-1 | Translated BLASTn Search Result | No |
| *Apis mellifera* | Bee | NP 001011617.1 | 1-53 | Abaecin-1 | Translated BLASTn Search Result | No |
| *Bombus affinis* | Bee | XP 050578474.1 | 22-85 | Abaecin-1 | Translated BLASTn Search Result | No |
| *Bombus ignitus* | Bee | AAQ90411.1 | 1-58 | Abaecin-1 | Translated BLASTn Search Result | No |
| *Bombus impatiens* | Bee | XP 003491544.1 | 1-58 | Abaecin-1 | Translated BLASTn Search Result | No |
| *Bombus pyrosoma* | Bee | XP 043604743.1 | 45-116 | Abaecin-1 | Translated BLASTn Search Result | No |
| *Bombus terrestris* | Bee | XP 003394701.1 | 22-85 | Abaecin-1 | Translated BLASTn Search Result | No |
| *Bombus vancouverensis nearcticus* | Bee | XP 033205111.1 | 22-85 | Abaecin-1 | Translated BLASTn Search Result | No |
| *Ceratina calcarata* | Bee | XP 017876439.1 | 1-55 | Abaecin-1 | Translated BLASTn Search Result | No |
| *Eufriesea mexicana* | Bee | XP 017760992.1 | 1-51 | Abaecin-1 | Translated BLASTn Search Result | No |
| *Frieseomelitta varia* | Bee | XP 043524279.1 | 1-54 | Abaecin-1 | Translated BLASTn Search Result | No |
| *Habropoda laboriosa* | Bee | XP 017790090.1 | 1-56 | Abaecin-1 | Translated BLASTn Search Result | No |
| *Heterotrigona itama* | Bee | CAD1479620.1 | 1-61 | Abaecin-1 | Translated BLASTn Search Result | No |
| *Melipona bicolor* | Bee | KAK1128096.1 | 1-50 | Abaecin-1 | Translated BLASTn Search Result | No |
| *Melipona quadrifasciata* | Bee | KOX74422.1 | 1-52 | Abaecin-1 | Translated BLASTn Search Result | No |
| *Athalia rosae* | Sawfly | XP 012260680.2 | 1-51 | Abaecin-1 | Translated BLASTn Search Result | No |

| | | | | | | |
|---|---|---|---|---|---|---|
| *Belonocnema kinseyi* | Wasp | XP 033225186.1 | 2-67 | Abaecin-1 | Translated BLASTn Search Result | No |
| *Copidosoma floridanum* | Wasp | XP 014208617.1 | 16-49 | Abaecin-1 | Translated BLASTn Search Result | No |
| *Fopius arisanus* | Wasp | XP 011309145.1 | 1-55 | Abaecin-1 | Translated BLASTn Search Result | No |
| *Leptopilina boulardi* | Wasp | XP 051165230.1 | 1-54 | Abaecin-1 | Translated BLASTn Search Result | No |
| *Leptopilina heterotoma* | Wasp | XP 043474498.1 | 1-53 | Abaecin-1 | Translated BLASTn Search Result | No |
| *Nasonia vitripennis* | Wasp | XP 003425826.1 | 1-59 | Abaecin-1 | Translated BLASTn Search Result | No |
| *Phymastichus coffea* | Wasp | XP 058790769.1 | 15-55 | Abaecin-1 | Translated BLASTn Search Result | No |
| *Pteromalus puparum* | Wasp | ABS44869.1 | 1-51 | Abaecin-1 | Translated BLASTn Search Result | No |
| *Trichomalopsis sarcophagae* | Wasp | OXU21302.1 | 34-67 | Abaecin-1 | Translated BLASTn Search Result | No |
| *Vespa crabro* | Wasp | XP 046820244.1 | 116-184 | Abaecin-1 | Translated BLASTn Search Result | No |
| *Vespula germanica* | Wasp | KAF7405906.1 | 1-56 | Abaecin-1 | Translated BLASTn Search Result | No |
| *Vespula pensylvanica* | Wasp | KAF7429375.1 | 55-117 | Abaecin-1 | Translated BLASTn Search Result | No |
| *Vespula vulgaris* | Wasp | KAF7402542.1 | 55-120 | Abaecin-1 | Translated BLASTn Search Result | No |
| *Atta texana* | Ant | QEPB01039057.1 | 6713-6811 | Abaecin-2 | Translated BLASTn Search Result | No |
| *Crematogaster levior* | Ant | CABEGB010000914.1 | 269983 -270081 | Abaecin-2 | Translated BLASTn Search Result | No |
| *Temnothorax nylanderi* | Ant | JASTWM010000507.1 | 4649-4747 | Abaecin-2 | Translated BLASTn Search Result | No |
| *Acromyrmex charruanus* | Ant | JAANIC010005180.1 | 528137 -528235 | Abaecin-2 | Translated BLASTn Search Result | No |
| *Acromyrmex echinatior* | Ant | XP 011049187.1 | 1-54 | Abaecin-2 | PSI-BLAST Search Result | Yes |
| *Acromyrmex echinatior* | Ant | JANHNX010000014.1 | 2396933-2397031 | Abaecin-2 | Translated BLASTn Search Result | Yes |
| *Acromyrmex heyeri* | Ant | JAANIB010010238.1 | 325671-325769 | Abaecin-2 | Translated BLASTn Search Result | No |
| *Acromyrmex insinuator* | Ant | JAANHZ010000870.1 | 103064 -103162 | Abaecin-2 | Translated BLASTn Search Result | No |
| *Anoplolepis gracilipes* | Ant | JAOBUE010000016.1 | 3450394 -3450555 | Abaecin-2 | Translated BLASTn Search Result | No |
| *Aphaenogaster ashmeadi* | Ant | NJRQ01000102.1 | 140396-140575 | Abaecin-2 | Translated BLASTn Search Result | No |
| *Aphaenogaster floridana* | Ant | NJRP01000116.1 | 328977-329201 | Abaecin-2 | Translated BLASTn Search Result | No |
| *Aphaenogaster fulva* | Ant | NJRO01000023.1 | 759564-759788 | Abaecin-2 | Translated BLASTn Search Result | No |
| *Aphaenogaster miamiana* | Ant | NJRN01000078.1 | 355578-355802 | Abaecin-2 | Translated BLASTn Search Result | No |
| *Aphaenogaster picea* | Ant | NJRM01000023.1 | 444152-444376 | Abaecin-2 | Translated BLASTn Search Result | No |
| *Aphaenogaster rudis* | Ant | NJRK01000139.1 | 123953-124177 | Abaecin-2 | Translated BLASTn Search Result | No |
| *Atta cephalotes* | Ant | ADTU01009315.1 | 9729-9827 | Abaecin-2 | Translated BLASTn Search Result | No |
| *Atta colombica* | Ant | LKEW01028451.1 | 1134 -1286 | Abaecin-2 | Translated BLASTn Search Result | No |
| *Atta colombica* | Ant | KYM77084.1 | 1-43 | Abaecin-2 | PSI-BLAST Search Result | No |
| *Camponotus floridanus* | Ant | AEAB01017815.1 | 1092-1181 | Abaecin-2 | Translated BLASTn Search Result | No |

| | | | | | | |
|---|---|---|---|---|---|---|
| *Camponotus pennsylvanicus* | Ant | JALHBL010000095.1 | 290527-290679 | Abaecin-2 | Translated BLASTn Search Result | No |
| *Camponotus vicinus* | Ant | JANXEZ010000009.1 | 13828362 -13828514 | Abaecin-2 | Translated BLASTn Search Result | No |
| *Cardiocondyla obscurior* | Ant | JADYXP010000024.1 | 2687357-2687431 | Abaecin-2 | Translated BLASTn Search Result | No |
| *Cataglyphis hispanica* | Ant | SGAY01002453.1 | 10768-10866 | Abaecin-2 | Translated BLASTn Search Result | No |
| *Cataglyphis niger* | Ant | SJPC01000773.1 | 43935-44033 | Abaecin-2 | Translated BLASTn Search Result | No |
| *Cyphomyrmex costatus* | Ant | LKEX01018418.1 | 22683-22781 | Abaecin-2 | Translated BLASTn Search Result | No |
| *Cyphomyrmex costatus* | Ant | KYM99867.1 | 5-75 | Abaecin-2 | PSI-BLAST Search Result | No |
| *Formica aquilonia x Formica polyctena* | Ant | CAJQTV010000019.1 | 337959-338093 | Abaecin-2 | Translated BLASTn Search Result | No |
| *Formica exsecta* | Ant | NPMM01010057.1 | 4705229-4705363 | Abaecin-2 | Translated BLASTn Search Result | No |
| *Formica selysi* | Ant | WHNR01000001.1 | 7214610-7214744 | Abaecin-2 | Translated BLASTn Search Result | No |
| *Harpagoxenus sublaevis* | Ant | JASTWR010000243.1 | 232292-232390 | Abaecin-2 | Translated BLASTn Search Result | No |
| *Lasius fuliginosus* | Ant | CASCKF010000903.1 | 58219-58356 | Abaecin-2 | Translated BLASTn Search Result | No |
| *Lasius niger* | Ant | LBMM01010681.1 | 809-925 | Abaecin-2 | Translated BLASTn Search Result | No |
| *Monomorium pharaonis* | Ant | BBSX02005052.1 | 8724-8834 | Abaecin-2 | Translated BLASTn Search Result | No |
| *Mycetomoellerius zeteki* | Ant | LKFA01009785.1 | 2530-2628 | Abaecin-2 | Translated BLASTn Search Result | Yes |
| *Mycetomoellerius zeteki* | Ant | XP 018307440.1 | 1-54 | Abaecin-2 | PSI-BLAST Search Result | Yes |
| *Odontomachus brunneus* | Ant | JAAGEC010019914.1 | 6890-7054 | Abaecin-2 | Translated BLASTn Search Result | No |
| *Pararachymyrmex cornetzi* | Ant | LKEY01038114.1 | 13348-13446 | Abaecin-2 | Translated BLASTn Search Result | No |
| *Paratrachymyrmex cornetzi* | Ant | KYN17872.1 | 1-54 | Abaecin-2 | PSI-BLAST Search Result | No |
| *Pogonomyrmex barbatus* | Ant | ADIH01028606.1 | 20225-20383 | Abaecin-2 | Translated BLASTn Search Result | Yes |
| *Pogonomyrmex barbatus* | Ant | XP 025074442.1 | 1-76 | Abaecin-2 | PSI-BLAST Search Result | Yes |
| *Pogonomyrmex californicus* | Ant | JAEVLP010000124.1 | 241705-241866 | Abaecin-2 | Translated BLASTn Search Result | No |
| *Polyergus mexicanus* | Ant | JAUDSZ010000018.1 | 6269843-6269911 | Abaecin-2 | Translated BLASTn Search Result | No |
| *Prenolepis imparis* | Ant | JAUDTA010000006.1 | 1132366-1132497 | Abaecin-2 | Translated BLASTn Search Result | No |
| *Pseudoatta argentina* | Ant | JAANIA010000071.1 | 8910073-8910171 | Abaecin-2 | Translated BLASTn Search Result | No |
| *Pseudomyrmex concolor* | Ant | QVOB01009298.1 | 29470-29565 | Abaecin-2 | Translated BLASTn Search Result | No |
| *Pseudomyrmex cubaensis* | Ant | QVOA01005862.1 | 11586-11684 | Abaecin-2 | Translated BLASTn Search Result | No |
| *Pseudomyrmex dendroicus* | Ant | QVNZ01010503.1 | 54-152 | Abaecin-2 | Translated BLASTn Search Result | No |
| *Pseudomyrmex elongatus* | Ant | QVNY01022936.1 | 5180-5278 | Abaecin-2 | Translated BLASTn Search Result | No |
| *Pseudomyrmex flavicornis* | Ant | QVNW01005555.1 | 183-278 | Abaecin-2 | Translated BLASTn Search Result | No |
| *Pseudomyrmex gracilis* | Ant | XP 020280064.1 | 16-78 | Abaecin-2 | Translated BLASTn Search Result | Yes |
| *Pseudomyrmex gracilis* | Ant | JWHX01000040.1 | 522150-522248 | Abaecin-2 | PSI-BLAST Search Result | Yes |

| | | | | | | |
|---|---|---|---|---|---|---|
| *Pseudomyrmex pallidus* | Ant | QVNV01002352.1 | 85602-85697 | Abaecin-2 | Translated BLASTn Search Result | No |
| *Pseudomyrmex sp.* | Ant | QVNX01019664.1 | 13231-13329 | Abaecin-2 | Translated BLASTn Search Result | No |
| *Solenopsis invicta* | Ant | PRJNA49629 | - | Abaecin-2 | BioProject via Zhang and Zhu (2012) | No |
| *Temnothorax americanus* | Ant | JASTWO010000091.1 | 223668-223766 | Abaecin-2 | Translated BLASTn Search Result | No |
| *Temnothorax curvispinosus* | Ant | QBEX01004092.1 | 321524-321619 | Abaecin-2 | Translated BLASTn Search Result | No |
| *Temnothorax longispinosus* | Ant | QBLH01000329.1 | 217005-217100 | Abaecin-2 | Translated BLASTn Search Result | No |
| *Temnothorax ravouxi* | Ant | JASTWP010000268.1 | 128869-128967 | Abaecin-2 | Translated BLASTn Search Result | No |
| *Temnothorax unifasciatus* | Ant | JASTWK010000092.1 | 127904-127999 | Abaecin-2 | Translated BLASTn Search Result | No |
| *Tetramorium alpestre* | Ant | JAJVSH010000150.1 | 315640-315738 | Abaecin-2 | Translated BLASTn Search Result | No |
| *Tetramorium bicarinatum* | Ant | VBVP01000517.1 | 25868-25966 | Abaecin-2 | Translated BLASTn Search Result | No |
| *Tetramorium immigrans* | Ant | VBVO01002714.1 | 11986-12084 | Abaecin-2 | Translated BLASTn Search Result | No |
| *Tetramorium parvispinum* | Ant | VBVR01000103.1 | 5246-5344 | Abaecin-2 | Translated BLASTn Search Result | No |
| *Tetramorium simillimum* | Ant | VBVQ01004268.1 | 8266-8364 | Abaecin-2 | Translated BLASTn Search Result | No |
| *Trachymyrmex septentrionalis* | Ant | LKEZ01022966.1 | 6697-6765 | Abaecin-2 | Translated BLASTn Search Result | No |
| *Vollenhovia emeryi* | Ant | BBUO01005684.1 | 5647-5775 | Abaecin-2 | Translated BLASTn Search Result | No |
| *Wasmannia auropunctata* | Ant | BBSV01098925.1 | 3288-3386 | Abaecin-2 | Translated BLASTn Search Result | Yes |
| *Wasmannia auropunctata* | Ant | XP 011698602.1 | 1-54 | Abaecin-2 | PSI-BLAST Search Result | Yes |
| *Mischocyttarus mexicanus* | Wasp | KAI4503845.1 | 1-49 | Abaecin-2 | Translated BLASTn Search Result | No |
| *Vespula germanica* | Wasp | KAF7405905.1 | 59-112 | Abaecin-2 | Translated BLASTn Search Result | No |
| *Vespula pensylvanica* | Wasp | KAF7429374.1 | 1-52 | Abaecin-2 | Translated BLASTn Search Result | No |
| *Vespula vulgaris* | Wasp | KAF7402541.1 | 1-49 | Abaecin-2 | Translated BLASTn Search Result | No |
| *Solenopsis invicta* | Ant | AEAQ02065247.1 | 1297704-1297841 | Abaecin-2 | Translated BLASTn search result | No |

* Sequences from chromosomes may not include sequence from the shorter second exon, as these are not sufficient to generate BLAST hits

**Table S3. ICP analysis sample metadata**

| | | | | | Number of samples used in ICP analysis | | |
|---|---|---|---|---|---|---|---|
| **Colony** | **Species** | **Collection area** | **Colony location (latitude, longitude)** | **Date collected** | **Worker ants** | **Fungus garden** | **Soil** |
| JKH000424 | *Atta texana* | Clear Creek Wildlife Management Area, Louisiana | (31.051, -93.401)* | 2021-05-19 | 1 | 0 | 0 |
| JKH000435 | *Atta texana* | Clear Creek Wildlife Management Area, Louisiana | (31.05082, -93.40154) | 2021-05-22 | 1 | 2 | 2 |
| JKH000423 | *Trachymyrmex septentrionalis* | Clear Creek Wildlife Management Area, Louisiana | (31.051, -93.401)* | 2021-05-19 | 1 | 1 | 2 |
| JKH000425 | *Trachymyrmex septentrionalis* | Alexandria State Forest, Louisiana | (31.11285, -92.4619) | 2021-05-21 | 1 | 1 | 2 |
| JKH000426 | *Trachymyrmex septentrionalis* | Alexandria State Forest, Louisiana | (31.11294, -92.46684) | 2021-05-21 | 1 | 1 | 3 |
| JKH000427 | *Trachymyrmex septentrionalis* | Alexandria State Forest, Louisiana | (31.11302, -92.46691) | 2021-05-21 | 1 | 0 | 2 |
| JKH000428 | *Trachymyrmex septentrionalis* | Alexandria State Forest, Louisiana | (31.11287, -92.4668) | 2021-05-21 | 1 | 2 | 1 |
| JKH000429 | *Trachymyrmex septentrionalis* | Alexandria State Forest, Louisiana | (31.11262, -92.46777) | 2021-05-21 | 1 | 0 | 1 |
| JKH000430 | *Trachymyrmex septentrionalis* | Alexandria State Forest, Louisiana | (31.11292, -92.46778) | 2021-05-21 | 1 | 1 | 2 |
| JKH000431 | *Trachymyrmex septentrionalis* | Alexandria State Forest, Louisiana | (31.11303, -92.46766) | 2021-05-21 | 1 | 0 | 2 |
| JKH000432 | *Trachymyrmex septentrionalis* | Alexandria State Forest, Louisiana | (31.11284, -92.46739) | 2021-05-21 | 1 | 1 | 2 |

| | | | | | | | |
|---|---|---|---|---|---|---|---|
| JKH000433 | *Trachymyrmex septentrionalis* | Alexandria State Forest, Louisiana | (31.11262, -92.46718) | 2021-05-21 | 1 | 0 | 2 |
| JKH000434 | *Trachymyrmex septentrionalis* | Alexandria State Forest, Louisiana | (31.11275, -92.46721) | 2021-05-21 | 1 | 1 | 2 |

* Provided coordinates are estimates

**Table S4. Susceptibility of *E. coli* JW 0473-3 (Δ*copA*) to *A. texana* and *A. colombica* abaecin-2 variants in combination with the pore-forming peptide cecropin A.** All assays were conducted in M9 media.

| | ***E. coli* JW 0473-3 (Δ*copA*)** | | | | |
|---|---|---|---|---|---|
| | $MIC_A$ | $MIC_B$ | $MIC_{combo}$ | FIC Index | Interpretation |
| Abaecin-2 WT AT + CecA | >128 | 1 | 8/0.125 | 0.31 | Synergy |
| Abaecin-2 H3A AT + CecA | >128 | 1 | 8/0.125 | 0.31 | Synergy |
| Abaecin-2 WT AC + CecA | >128 | 1 | 16/0.25 | 0.31 | Synergy |
| Abaecin-2 H3A AC + CecA | >128 | 1 | 8/0.125 | 0.31 | Synergy |

**Supplementary code S1.** Sample DynaFit scripts used for Cu(II) binding affinity determination of abaecin-2 peptides.

**A.** Sample script for the competition between WT abaecin-2 (12.8 µM) and DP4 (2 µM), assuming 1:1 binding stoichiometry to each abaecin-2 and DP4.

```
; M = Cu
; L = DP2, DP3, or DP4
; P = WT or H3A peptide

[task]
task = fit
data = equilibria

[mechanism]
M + L <==> ML     :  KdL  dissociation
M + P <==> MP     :  KdP  dissociation

[constants] micromolar
KdL = 7.94328e-9
KdP = 1e-7?

[concentrations] micromolar
L = 2
P = 12.8

[responses] F550norm
L = 0.5

[data]
variable     M
set          Data

[output]
directory ./WT+DP4/

[set:Data]
M     F550norm
```

**B.** Sample script for the competition between WT abaecin-2 (3.6 µM) and DP3 (4 µM), assuming 1:1 binding stoichiometry to each abaecin-2 and DP3.

```
; M = Cu
; L = DP3
; P = WT abaecin-2

[task]
```

```
task = fit
data = equilibria

[mechanism]
M + L <==> ML    :  KdL  dissociation
M + P <==> MP    :  KdP  dissociation

[constants] micromolar
KdL = 5.01187e-7
KdP = 1e-6?

[concentrations] micromolar
L = 4
P = 3.6

[responses] F550
L  = 0.25

[data]
variable   M
set        Data

[output]
directory ./WT+DP3/

[set:Data]
M     F550nm
```

**C.** Sample script for the competition between H3A abaecin-2 (12.8 µM) and DP2 (2 µM), assuming 1:1 binding stoichiometry to each abaecin-2 and DP2.

```
; M = Cu
; L = DP2
; P = H3A abaecin-2

[task]
task = fit
data = equilibria

[mechanism]
M + L <==> ML    :  KdL  dissociation
M + P <==> MP    :  KdP  dissociation

[constants] micromolar
KdL = 7.94328E-5
KdP = 1e-5?
```

```
[concentrations] micromolar
L = 2
P = 12.8

[responses] F550
L  = 0.5

[data]
variable   M
set        Data

[output]
directory ./H3A+DP2/

[set:Data]
M     F550nm
```
